# Cell-Dense Bioink Design for Xolography: Coupling Refractive Index-Matching with Increased Photoreactivity

**DOI:** 10.64898/2026.06.03.729865

**Authors:** Aiste Balciunaite, Sebastian Inacker, Asia Badolato, Erik Brauer, Niklas Felix König, Lucíola Vasconcelos Lima, Grace Rachel Humphreys, Claudiadele Polinari, Samuel Palato, Pablo Paniagua Hernandez, Miriam Filippi, Stefan Hecht, Robert K. Katzschmann

## Abstract

Bioxolography enables high-resolution fabrication of geometrically complex, cell-laden constructs for tissue engineering. However, tissue-relevant cell densities conflict with the optical transparency required for efficient dual-color volumetric printing. In this work, we extend the Bioxolography toolbox to include refractive index (RI) matching for cell-laden bioresins using iodixanol (IDX). Remarkably, IDX enhances optical transparency and boosts reactivity — a phenomenon unique to Xolography. Yet, excessive IDX compromises dual-color efficiency through increased absorption and undesired UV-only curing, underscoring a central trade-off between optical clarity and photochemical performance. Systematic tuning of resin compositions along an iso-refractive index line demonstrated the versatility of Bioxolography, with IDX enhancing polymerization and 4-Hydroxy-TEMPO providing biocompatible inhibition. Optimizing composition and printing parameters yielded GelMA hydrogels with cell densities up to 5·10^6^ cells·mL^−1^. Cell-laden prints achieved sub-100 µm resolution and complex geometries such as channels and gyroids. Using skeletal muscle tissue as a model, we validated RI matched Bioxolography as a promising strategy for tissue engineering by demonstrating cell alignment along printed grooves and formation of mature muscle fibers characterized by MyoHC+ staining and fusion index. By integrating physical, chemical, and biological perspectives, this work advances Xolography toward biomaterials development and reinforces its position as an emerging volumetric (bio)printing technology.

**Table of Contents:** 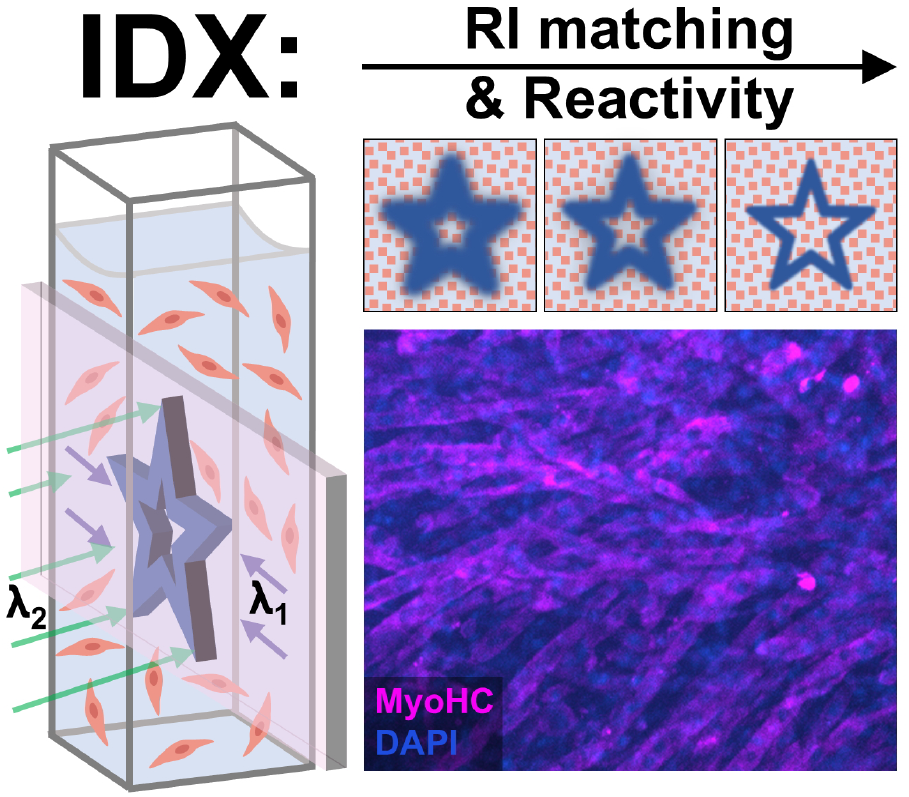

For printing higher cell density bioresins with Xolography, iodixanol (IDX) is added for refractive index-matching. The addition leads to an unexpected additional effect with increased reactivity in the dual-color photopolymerization. With careful adjustment of the resin composition and the printing parameters, Bioxolography is proven as a viable tool for tissue engineering.

## Introduction

Volumetric 3D printing techniques represent the frontier of light-based additive manufacturing. By printing objects directly within the volume of a resin, these methods allow for significantly faster fabrication time, larger geometric freedom of design, and high resolution.^1^ These advantages are particularly relevant for tissue engineering applications, where both macro- and micro-scale architectures are crucial for supporting tissue development and function.^2^ As such, volumetric bioprinting (VBP) is a promising biofabrication approach for creating advanced 3D cell culture systems with diverse cell types, where cell signaling and cell-to-cell contacts drive the tissue morphogenesis process.^3,4^ Yet, light-based printing techniques favor minimizing the number of cells in the resin as they act as scattering centers, disrupting light propagation and reducing printing resolution. Refractive index matching of the surrounding matrix to the cells, using compounds such as iodixanol (IDX), can increase transparency by reducing scattering.^5,6^ In addition to its high refractive index (RI) due to the large number of heteroatoms, IDX is cytocompatible and chemically inert for light-based volumetric printing at 405 nm.^5–8^ In tomographic VBP, IDX has been used to optically tune bioresins and print mm-scale functional perfusable structures with liver organoids.^5^ However, these organoids remain as isolated functional units rather than forming a cohesive tissue. Filamented light (FLight) bioprinting was developed as an alternative light-based approach to fabricate highly aligned constructs with densely packed cells.^7,9,10^ FLight-printed constructs have uniaxially formed microbeams that provide microtopographical cues for cell alignment and tissue formation. However, the technique has inherent limitations in multidirectionality and microbeam resolution at higher cell densities.^7^ The addition of refractive index matching substances in FLight printing results in reduced formation of microbeams, as it prevents the RI mismatch between polymerized and unpolymerized resin required for the self-focusing effect.^8^ Therefore, printing methods that combine high resolution in three dimensions with the ability to support high cell densities are not available yet.

A recently developed volumetric printing technique, Xolography, leverages dual-color photoinitiators (DCPI) to spatially control polymerization at the intersection of a UV light sheet and a visible light projection.^11^ Upon exposure to light of the first wavelength λ_1_ (375 nm) in the UV light sheet, the DCPI molecules switch from their dormant form to an active isomer, which reverts thermally. Due to the associated changes in the molecular absorbance, the active form can be excited orthogonally with visible light of the second wavelength λ_2_ (500-650 nm). Dual excitation by both colors initiates the polymerization cascade in the presence of suitable coinitiators. By sweeping the UV light sheet through the volume with continuous updating of the visible light projection, the object is materialized inside the volume.^11^ Unlike the threshold-based control used in tomographic volumetric printing, the AND-logic mechanism of Xolography DCPIs enables binary control over polymerization, eliminating inherent non-linearities and offering an alternative method to achieving high flexibility in shape design. While the first demonstration of Xolography was limited to highly crosslinked hydrophobic photopolymers, recent works have focused on the expansion of Xolography to more complex material formulations, such as organic multi-materials or organic-inorganic hybrid materials, as well as hydrogel materials^12–18^

Bioxolography, the application of Xolography to tissue engineering, connects the inherent high resolution and geometric complexity of Xolography with cell compatibility – features critical for fabricating tissue constructs with fine microarchitecture.^2,19–21^ Initial work by Stoecker et al. demonstrated xolographic printing of hydrogel-based constructs containing small cell aggregates using a biocompatible coinitiator (BisTris, BT), though cell viability was limited to regions with minimal resin exposure.^22^ Subsequent advances through tuning of the resin composition have improved cytocompatibility to more than 80%.^23,24^ Nonetheless, these efforts were limited to low cell densities up to 1·10^6^ cells·mL^-1^, far below the high initial seeding densities typically required for robust tissue formation, preventing the use of Bioxolography for tissue engineering applications.

In this work, we introduce a material design strategy for refractive index-matched Bioxolography to print constructs with tissue-relevant cell densities (≥ 5·10^6^ cells·mL^-1^ for skeletal muscle), previously inaccessible with conventional xolographic approaches.^4,20,21^ We identify iodixanol (IDX) as a dual-function additive that not only suppresses light scattering in dense cell suspensions, markedly improving print fidelity, but also unexpectedly enhances polymerization efficiency — a previously unreported effect specific to dual-color photoinitiator (DCPI) systems, and never described for single-color photoinitiators.

In direct adaptation of the Xolography workflow (Figure 1), we investigated the underlying optical and photochemical relations between the components to develop cell-dense bioresins with methacrylated gelatin (GelMA) and murine myoblasts (C2C12) for skeletal muscle tissue engineering. Guided by mechanistic investigations of the chemical processes involving the DCPI, coinitiator, and IDX, bioresin compositions were tuned for reactivity along an iso-refractive index line using IDX as a polymerization booster and 4-Hydroxy-TEMPO (TEMPOL) as antagonistic radical polymerization inhibitor. With further printing parameter optimization, this strategy allowed high-resolution printing of constructs with an initial cell seeding density of 5·10^6^ cells·mL^-1^ achieving feature sizes down to 70 µm and freestanding structures of 100 µm. Importantly, the printed constructs supported tissue development, as demonstrated by cell alignment along printed micro-structures, cell differentiation, and mature myotube formation. Stemming from an in-depth analysis of the physical, chemical, and biological interactions involved, we introduce iodixanol as a dual-function additive for designing functional (bio)materials for Xolography. This work lays the foundation for Bioxolography as a powerful tool for tissue engineering, *in vitro* models, and regenerative medicine for biofabrication of biomimetic tissues.

**Figure 1.**
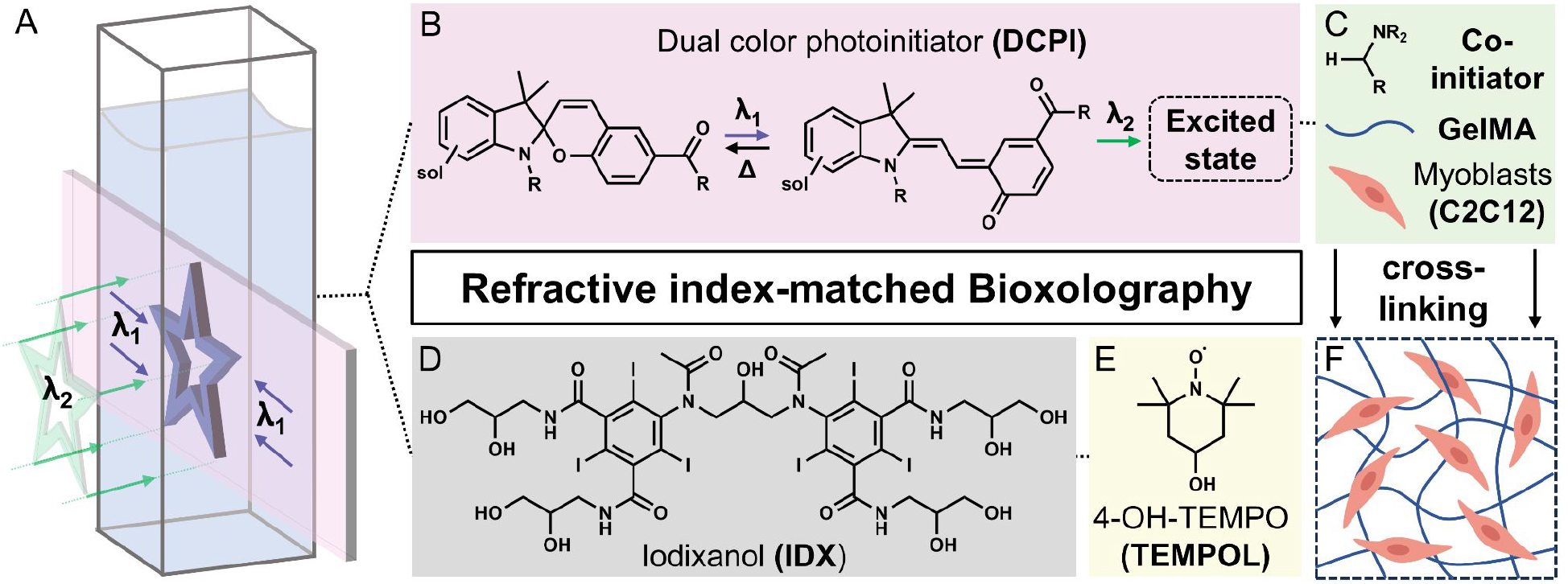
Schematic illustration of refractive index-matched Bioxolography with key steps and components. (A) Xolographic printing setup: the printing mixture is illuminated from orthogonal directions by a UV light sheet (λ_1_) and a visible light projection (λ_2_). Polymerization of the desired object occurs at the intersection of the two light beams with different wavelengths. (B) Photochromism of the dual-color photoinitiator (DCPI, sol = water-solubilizing group): exposure to λ_1_ activates the DCPI, subsequent λ_2_ illumination yields an excited state that can react with the coinitiator (C) to initiate crosslinking of the GelMA-cell suspension. Iodixanol (D, refractive index-matching substance) and TEMPOL (E, radical polymerization inhibitor) optimize printing resolution in Bioxolography volumetric 3D printing (F) in the presence of high myoblast concentrations (5·10^6^ mL^-1^).

## Results and Discussion

### Refractive index-matching under dual-color conditions

We first addressed the problem of light scattering due to the presence of cells in the printing resin by considering the use of Iodixanol (IDX) in Xolography as a biocompatible, non-toxic, and iso-osmotic RI matching additive, in equivalence to tomographic VBP studies using 405 nm light.^5^ Unlike single-color VBP techniques, Xolography relies on the intersection of a UV light sheet and a visible light projection. Hence, enabling high-cell-density Xolography requires maintaining optical homogeneity at both wavelengths — a more complex requirement than in 405 nm single-beam systems. To investigate this, we designed a custom-built setup to measure transmittance and assess light scattering and absorption in cell-laden mixtures under dual-wavelength conditions (Figure S1). The system measures transmittance loss in the UV (375 nm) and visible (565 nm) regions, matching the absorbance spectra of DCPI5004 and the emission profile of the Xube^2^ volumetric printer. We used this setup to evaluate the effect of an increasing IDX concentration in the mixture. We hereby measured the degree of turbidity for C2C12 myoblast dispersions in a GelMA matrix (10% w/v in water, 186 mM BT) at a cell concentration of 5·10^6^ mL^-1^. The results are summarized in Figure 2A.

**Figure 2.**
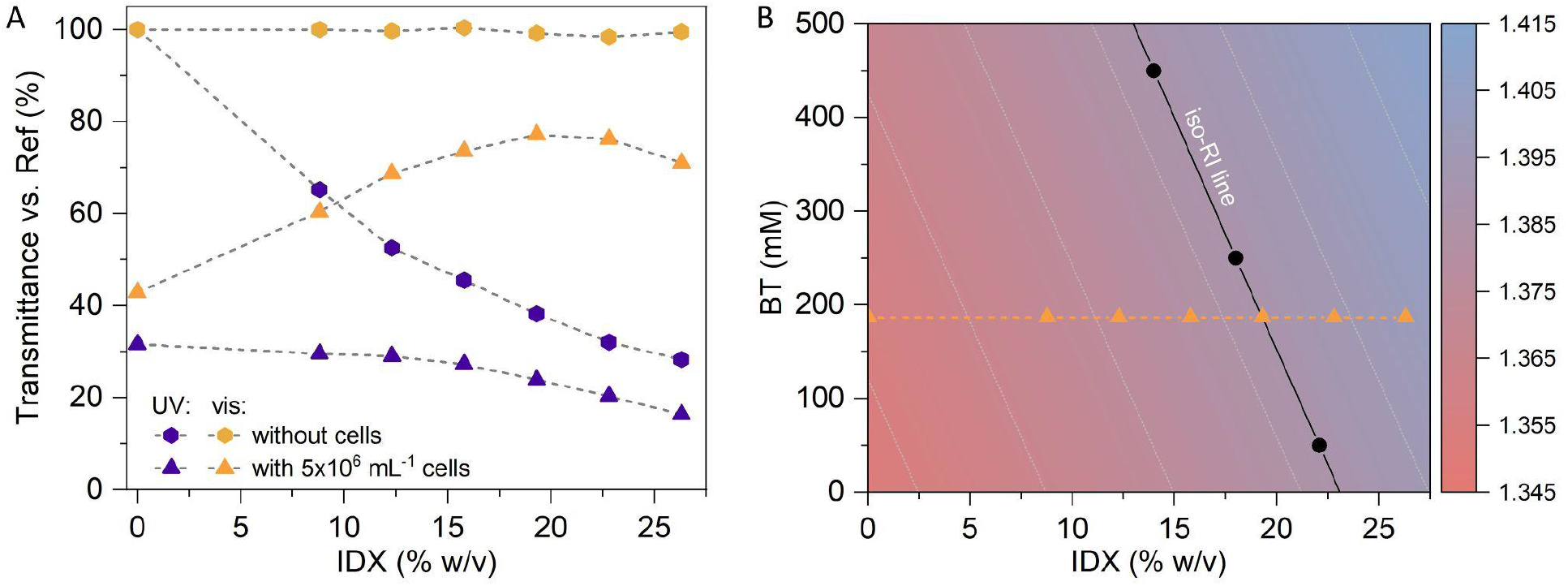
Optical characterization of cell dispersions in GelMA matrix (186 mM BT) at both printing wavelengths. (A) Determined transmittance for each cell composition for UV and visible spectral ranges (central wavelengths: 375 nm and 565 nm respectively, cuvette path length 1 cm). With an increasing IDX concentration, the visible light transmittance increases, whereas it is strongly reduced for the UV wavelength. (B) Refractive index map for different IDX and BT concentrations in GelMA with data from A included (orange triangles) and iso-RI line at 1.388. Representative compositions along these lines are marked with black dots: 450 mM BT / 14.0% w/v IDX, 250 mM BT / 18.0% w/v IDX and 50 mM BT / 22.1% w/v IDX.

The measured degree of transmittance for the visible light regime increased with an increasing IDX concentration from ∼43% to ∼77% at 19.3% w/v IDX, suggesting that the RI of the medium approaches that of the primary scattering structures within the cells, minimizing contrast and thus scattering. However, at higher IDX concentrations, transmittance decreased again — likely due to overcompensation, where the medium becomes optically denser than the cells, reintroducing RI mismatch and enhancing scattering. In contrast, while iodixanol addition increases visible light transmittance, it also raises UV absorbance, making its use a trade-off that must be optimized for effective dual-color printing. For the UV region around 375 nm, the transmittance decreased with additional IDX with and without cells. Although the extinction coefficient of IDX in water at 375 nm is small (3.20 L·mol^-1^·cm^-1^, respective 2.1 (% w/v)^-1^·cm^-1^), dissolving a 18% w/v solution results in a reduced transmittance of <40%, (OD = 0.4, Figure S3) at the light sheet wavelength. This higher absorbance results in a strong light sheet intensity attenuation toward the center of the cuvette.

While this effect is partially balanced by the printer hardware as the UV light sheet is built-up from both sides of the cuvette, it is still important to ensure homogeneity of the UV light sheet over the complete width of the cuvette.^11^ Otherwise, these inhomogeneities directly translate to different concentrations of the active form of the DCPI in the light sheet plane. After the second excitation, these differences result in a lower degree of initiation and therefore a lower degree of conversion at the center of the printing vat (see Supporting Information, Figure S4 for a more detailed discussion). Notably, the extinction coefficient for 405 nm is ∼20 times smaller (0.16 L·mol^-1^·cm^-1^, respective 0.10% w/v^-1^·cm^-1^, Figure S2), leading to a higher intrinsic transparency at the commonly used 405 nm wavelength for other volumetric 3D printing techniques such as tomographic volumetric printing or FLight.

Based on the dual-color illumination, the optical properties must be carefully optimized at both wavelengths, as the required RI-matching depends on both cell content and matrix composition, given the intrinsic optical heterogeneity of biological cells.^6,7,25–27^ Reference experiments in aqueous cell suspensions (Figure S2) showed that increasing cell density requires higher IDX concentrations to approach refractive index matching; however, even under matched conditions, substantial visible-light scattering and transmission losses persist at higher cell densities. The associated increase in UV absorbance by iodixanol strongly attenuates the UV light sheet, limiting dual-color activation and thereby defining a practical upper limit for the achievable cell density in the present Xolography printing system.

To address the issue of increasing OD at higher IDX concentrations, we aimed to reduce the amount of IDX. Measuring the RI in dilution series of aqueous solutions revealed that higher concentrations of BT can also modulate the RI of the GelMA matrix at the relevant visible light wavelength regime. The resulting RI is a linear combination of the contributions from IDX and BT, enabling the construction of a RI map (Figure 2B). This linearity allows for the prediction of the overall RI based on the formulation composition. By integrating the transmittance measurements shown in Figure 2A, we identified an optimal RI value of 1.388 at the used concentrations of 19.3% w/v IDX / 186 mM BT for minimizing scattering in the cell-laden samples. This value is in accordance with literature-reported cell RI values, providing a rational basis for its selection as the target RI.^26,27^ Along the iso-RI line at 1.388, several formulations were identified that achieved equivalent RI values with varying IDX/BT ratios; for example, 14.0% w/v IDX / 450 mM BT and 22.1% w/v IDX / 50 mM BT. Despite their matched optical properties, these formulations likely exhibit rather different photopolymerization reactivities, as the efficiency in Xolography critically depends, among other factors, on the coinitiator concentration.^15^

### Combining refractive index-matching and dual-color reactivity

To evaluate how different IDX/BT compositions affected the dual-color reactivity under iso-RI conditions, we quantified the dual color reactivity along the iso-RI line introduced in Figure 2 (see Supporting Information, Table S1 for full dataset). Gelation threshold times of aqueous poly(ethylene glycol) diacrylate (PEGDA) solutions were used as a reference system, enabling external monitoring of reactivity changes for defined combinations of DCPI, BT, and IDX. In addition, the OD at the light-sheet wavelength 375 nm was calculated from the IDX concentration to evaluate potential UV light sheet intensity attenuation and its effect on light sheet homogeneity. The reactivities were investigated both under UV+Vis as well as UV-only illumination to also account for potential unfeasible side reactions. To evaluate this selectivity, the ratio between UV+Vis and UV-only curing was quantified as the dual-color reactivity gain (DCRG).

A clear trend emerged: formulations with a lower IDX content but higher BT concentration exhibited higher overall photoreactivity with higher dual color reactivity gain, while displaying reduced OD_375 nm_ values. These factors promote an efficient and uniform printing process. Moreover, excessive IDX, though beneficial for RI matching, can limit dual-color efficiency due to increased absorption and undesired UV-only induced reaction. This highlights a central trade-off in resin formulation between optical clarity and photochemical performance. Importantly, the IDX-formulation without BT showed only moderate reactivity under dual-color conditions, reinforcing that IDX itself does not act as a coinitiator. Thus, BT is essential for effective visible-light-induced radical formation from the DCPI in the bimolecular activation process required for Xolography.

### Influence of IDX on dual-color photopolymerization

With this, we sought further insight into the mechanistic interplay of the molecular components by systematically investigating the role of IDX in the dual-color polymerization cascade. The presence of IDX increased the reactivity of the DCPI5004 system by at least a factor of 2.4 under the investigated conditions using the same PEGDA reference system (Table S2). While the DCPI–BT system without IDX exhibited a DCRG of 51.9%, the addition of IDX reduced this value to 31.0%, indicating an increased contribution of UV-only initiation despite the overall reactivity enhancement. IDX was further found to induce polymerization upon UV illumination even in the absence of DCPI. The system reactivity depends on the coinitiator presence, as BT-free mixtures required high UV intensities for curing. The absence of any difference between UV+Vis and UV-only curing (zero DCRG) in DCPI-free mixtures confirms that initiation proceeds exclusively via UV light in this case.

Spectroscopic experiments revealed the release of molecular iodine (I_2_) from IDX under intense UV illumination in aerobic solvents (Figure S5).^28^ From I_2_, iodine radicals are generated which act as initiating species even at low concentrations.^29^ The initiation ability was strongly enhanced in the presence of tertiary amines, as the generated iodine species introduce amine-based radical formation via a hydrogen abstraction mechanism.^30^ A related increase in radical-mediated reactivity has been reported for the addition of aryl iodides.^31^ Thus, we conclude that the described UV light-triggered iodine release from IDX and its complex influence in radical polymerization leads to non-specific crosslinking in the context of Xolography.^32–35^ Interestingly, no UV-only induced reaction takes place from IDX when illuminated with 405 nm light with the same intensity as for 375 nm, confirming that this side reaction does not occur in single-wavelength based techniques.^5–7^

Iodine release from IDX and its subsequent reactions with BT, however, cannot fully account for the enhanced reactivity of the type II DCPI photo cycle (Figure 4). Notably, the combination of all three components results in an even lower dual-color polymerization threshold, indicating that additional effects must be considered: the addition of IDX changes the mixtures’ polarity, leading to a slight bathochromic shift in the absorbance maximum of the active MC isomer (and increased thermal half-life (Figures S6 and S7).^15,36^ Additionally to bulk polarity changes, the local environment is affected by the presence of IDX. Considering the large excess of both the coinitiator and IDX relative to DCPI, the behavior of the photoexcited merocyanine form MC* following visible-light excitation is a key factor for the efficiency of the dual-color process. MC*-type motifs are known to exhibit complex excited-state dynamics arising from multiple metastable isomers with different central double-bond configurations, whose stabilization and interconversion barriers strongly depend on factors like solvation network, viscosity and hydrogen-bonding interactions.^37–39^ Changes in the local environment, e.g. in the microviscosity, can therefore directly affect MC* relaxation pathways and lifetimes, providing a plausible contribution to the enhanced reactivity observed in the presence of IDX.^40–43^ Consistent with this, time-resolved fluorescence measurements (Figure S8) show an accelerated decay of the emissive excited state in the presence of IDX, indicating changed relaxation pathways of MC*. Additionally, the effect of iodine-containing additives on dye photophysics is inherently complex, as they can concurrently promote intersystem crossing, alter excited state lifetimes, and participate in charge-transfer and redox processes, with outcomes that depend sensitively on the chromophore and excitation conditions.^44^ Blend experiments with iodixanol (IDX) and iohexol (IOX) at constant total iodine concentration (Figure S9) further demonstrate that the reactivity enhancement cannot be attributed to iodine content alone. While IOX increases reactivity relative to the iodine-free reference, it remains less effective than IDX, and increasing the IDX fraction leads to a gradual increase in reactivity. This behavior highlights the importance of the molecular identity of the iodinated additive and supports the presence of multiple, interdependent contributions, including local environment effects and scaffold-dependent photophysical interactions.

This complexity is further reflected in the formation of photoproducts during polymerization. An additional “colored species” is observed, evident from changes in sample appearance (Figure 4A), indicating altered reaction pathways in the presence of IDX. Absorbance measurements reveal that, in addition to the expected SP and MC states, a third species forms during dual-color polymerization, exhibiting a distinct band at 490–500 nm (Figure 4B,C). This species is attributed to a DCPI-derived photoproduct formed via reaction of excited MC with the coinitiator, yielding a hydroxy-functionalized radical that acts as a chain-terminating species.^11,45^ With the intact dye backbone, the formation of colored byproducts as a dead-end of the photoreaction is expected.^45,46^ In the presence of IDX, formation of this species is significantly accelerated under dual-color irradiation. Tracking the ratio of the absorbance of this species to the MC form (Figure 4D) shows a continuous increase in IDX-containing mixtures (purple), whereas IDX-free systems exhibit only delayed buildup (orange).

Together, these observations demonstrate that IDX influences both the kinetics and pathways of dual-color photopolymerization. However, further elucidation of these processes will be essential for a complete mechanistic understanding and motivates additional transient and steady-state spectroscopic studies, particularly in view of recent reports describing oxidative pathways in related DCPI systems.^24,42,47,48^

### Iso-RI composition printing performance and characterization

Our next goal was to translate these findings to a bioprintable resin. While the preceding experiments enabled extensive investigations into the molecular interplay using the PEGDA reference system, it also emerged that balancing all components (DCPI, BT, IDX and TEMPOL) is essential for achieving optimal resin performance.

Building on this three-component system, we tested the use of water-compatible radical polymerization inhibitors ascorbic acid (AA), pyrogallol (PG) and 4-Hydroxy-TEMPO (TEMPOL). The addition of inhibitors to enhance printability is well established in volumetric 3D printing.^13,49– 51^ In our case, the main intent for the inhibitor addition was to counteract the UV-only induced polymerization from IDX, thereby increasing printing contrast. Using the PEGDA reference system, the inhibition ability of all tested substances was confirmed with TEMPOL being the most potent one, both for the overall reactivity and the DCRG (Figure S10). Ascorbic acid and pyrogallol also reduced polymerization reactivity in a concentration-dependent manner, but to a much lower extent, in agreement with literature.^52^ In comparison, the use of TEMPOL introduced only minimal UV attenuation while maintaining effective inhibition, making 5 ppm the most favorable concentration for preserving dual-color performance.

We selected 10% w/v GelMA as the polymerizable matrix due to the widely established biocompatibility and prior use in Xolography.^5,22,24^ We additionally included fibrinogen (4 mg·mL^-1^) to promote cell spreading and viability.^53^ As outlined in Figure 2, we identified several candidate formulations containing different ratios of IDX and BT on the iso-RI line optimized for cell-laden resin optical transparency and then screened the mixtures for printing feasibility, resolution, and mechanical properties. The selected mixtures were: 450 mM BT and 14.0% w/v IDX (14IDX), 250 mM BT and 18.0% w/v IDX (18IDX), 50 mM BT and 22.1% w/v IDX (22IDX). A control mixture of 450 mM BT and without IDX (0IDX) was also tested. All formulations contained constant initiator and inhibitor concentrations at 300 ppm DCPI and 5 ppm TEMPOL, respectively.

We investigated the printing performance of the GelMA-based mixtures to understand the achievable resolution and structure complexity by tuning the parameters of UV intensity and light-sheet translational speed (Figure 5A) for each mixture. To investigate the achievable x-y resolution, we printed grids with positive and negative spacings of 50 µm to 300 µm (Figure 5B) and stars to quantify the positive and negative features (Figure 5C), following procedures reported for quantifying resolution in volumetric printing.^22,24,54^ The outer tips and the inner hollow tips of the stars were used to measure the positive resolution and negative resolution, respectively (Figure 5D and 5E). The values for 0IDX were highest (positive: 211 ± 98 µm; negative: 66 ± 23 µm), followed by 22IDX (85 ± 45 µm; 175 ± 57 µm), 18IDX (80 ± 30 µm; 110 ± 26 µm) and 14IDX (67 ± 35 µm; 70 ± 24 µm) (Figure 5F). For positive resolution, 0IDX differed significantly from all IDX-containing resins, with no significant differences found between the IDX-containing groups (Table S3). For negative resolution, no significant difference was observed between 0IDX and 14IDX, while all other comparisons were significant (Table S4). Interestingly, the addition of IDX to the printing samples resulted in a better positive resolution but a decrease in the achievable negative resolution at higher IDX concentrations, though this decrease was recovered in the 14IDX samples. This effect showcases the boosting of the intended dual-color reaction for vis light illuminated positions (positive features), but also reveals the IDX-mediated UV-only reaction detrimental for negative features along the UV-light sheet plane. In fact, the stars printing with the 22IDX mixture had a strong asymmetry (Figure S11) which is likely due to the high amount of IDX present in the solution. Overall, even in cell-free resins we concluded that IDX helps to increase the printing performance due to the synergistic effect of IDX on the DCPI.

Aside from differences in resolution, the different mixtures required different energies to form 3D structures. As indicated by the colorbar in Figure 5F, the structures for resolution analysis were printed at 2 mJ·mm^-2^ UV intenstiy for 14IDX, 5 mJ·mm^-2^ for 18IDX, 13 mJ·mm^-2^ for 22IDX, and 27 mJ·mm^-2^ for 0IDX. The overall reduction in energy required for IDX-containing mixtures compared to IDX-free systems is attributed to the enhanced reactivity of DCPI in the presence of IDX. However, within IDX-containing formulations, increasing IDX concentration leads to higher required energy doses. This is due to two factors: increased UV absorbance at higher IDX concentrations, which reduces the effective UV intensity, and a simultaneous decrease in BT concentration along the iso-refractive index line, which lowers photoreactivity (Figure 3A). Consequently, higher UV and relative visible light doses are required at higher IDX contents. Interestingly, we measured the storage modulus of all the mixtures and found no significant differences, showing that the changes in energy values required to print in the different mixtures did not have an effect on the mechanical properties of the gel (Figure S12).

**Figure 3.**
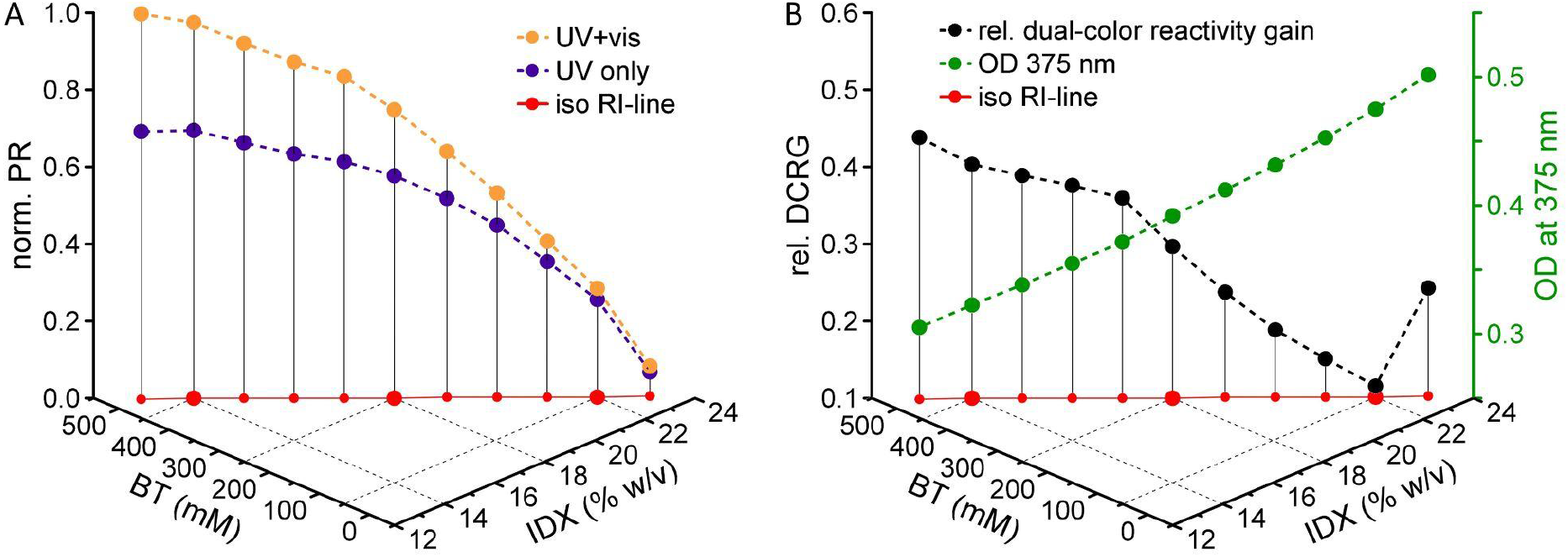
Optimizing dual-color photoreactivity and optical transparency. (A) Normalized photoreactivity for different BT/IDX concentrations along the iso-RI line (red, data taken from Figure 2). All compositions were examined under UV+vis (orange) and UV only (purple) conditions. (B) Calculated dual-color reactivity gain (black) based on the values in A, indicating the favored dual-color reaction over the UV-only induced reaction for lower IDX contents. Similarly, lower IDX contents lead to a reduced optical density at 375 nm (green), ensuring a higher light-sheet homogeneity.

We recorded complete printing windows of the 14IDX and 18IDX printing mixtures to define the thresholds for reliable 3D structure formation as a function of the UV intensity (energy, mJ·mm^-2^) and light sheet z-translation speed (mm·min^-1^) (Figure 5G). We used a resolution grid (Figure S13) and characterized the structures as undercured, well-printed, or overcured. At 23 °C, the required energies for printing the 14IDX mixtures increased from 4–8 mJ·mm^-2^ to 7–9 mJ·mm^-2^ for the 18IDX.

Importantly, we found that temperature control during gelation and printing was critical, as lower temperatures increased both the fraction of DCPI in its active MC (based on a larger t_1/2_) and the increased crosslinking rates of GelMA.^24^ The thermal half-life of the DCPI is strongly dependent on temperature, with a value of 4.7s at 16 °C compared to 2.7s at 25 °C (Figure 5H). We found that colder printing temperatures resulted in a decrease of the required printing energies for both mixtures: a decrease from 4–9 mJ·mm^-2^ (23°C) to 2–6 mJ·mm^-2^ (16°C) for the 14IDX mixture, and from 7–9 mJ·mm^-2^ (23°C) to 4–8 mJ·mm^-2^ (16°C) for the 18IDX mixture (Figure 5G). Therefore, we concluded that printing at lower temperatures was beneficial to minimize the UV exposure to biological samples, as lower UV energies were needed to obtain sufficient DCPI activation for structure formation. We also concluded that stable temperature control of the bioresin during gelation and printing was critical for reproducible printing conditions.

Based on the results of the x-y resolution and the printing windows, we chose to use the 14IDX resins for all further experiments at a printing temperature of 16 °C. To demonstrate the z-resolution, we printed a branching channel structure with channel diameters of 1 mm, 750 µm, and 500 µm (Figure 4I), and a gyroid with wall thickness of 1 mm (Figure 5J). We found that obtaining a finer z-resolution was more challenging than in x-y-direction, as the continuous visible light projection causes curing to continue even in segments where the light-sheet already passed through.^15^

**Figure 4.**
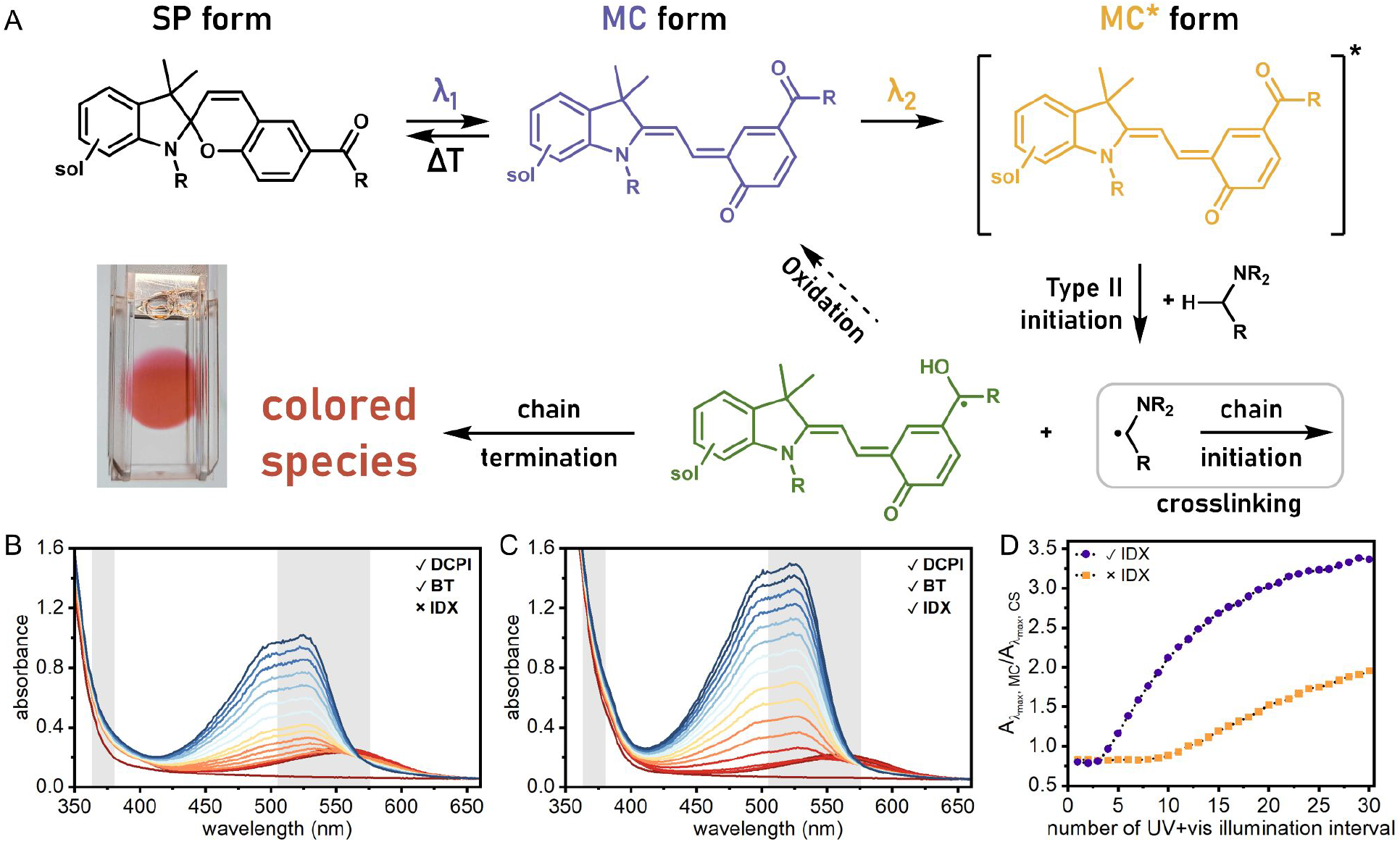
(A) Photoreaction cascade of the DCPI. Upon irradiation with λ_1_ (UV, 375 nm), the spiropyran form (SP, black) is isomerized to the open merocyanine form (MC, purple). Subsequent excitation with λ_2_ (visible, 565 nm) generates the excited state MC* (yellow) and introduces hydrogen abstraction from the amine-based coinitiator for starting radical formation. This reaction yields the reduced DCPI-centered benzhydrol radical (green), ultimately leading to the formation of an additional “colored species” from radical termination reactions. (B, C) This colored species was detected via UV–Vis absorbance and shows an absorbance band around 522 nm. (D) The formation of the colored species under dual-color illumination was monitored by tracking the ratio between absorbance at the DCPI λ_max_ (554 nm or 558 nm) and at the colored species λ_max_ (523 nm or 522 nm). In the presence of IDX (purple), the colored species appears more direct, as indicated by the steeper increase in this ratio compared to the IDX-free system (orange).

**Figure 5.**
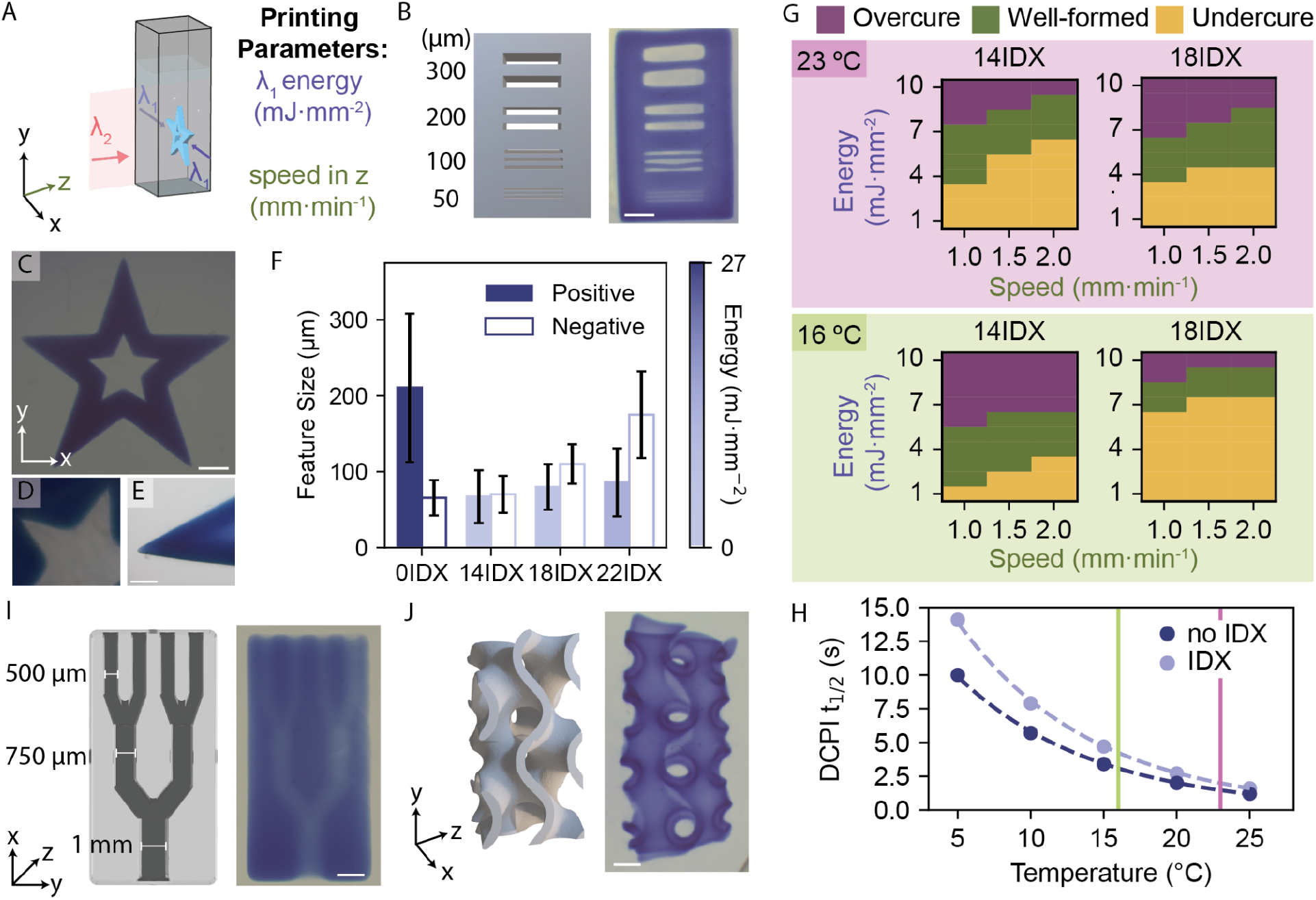
Characterization of bioresin printing behavior. (A) Definition of the axes and printing parameters in xolography. (B) Image of a grid resolution printed in 14IDX, with feature sizes of 300 µm, 200 µm, 100 µm, and 50 µm. (C) Star-shaped construct printed to analyze the positive and negative features. (D) Close-up of a positive pointed star. (E) Close-up of the negative features analyzed for resolution. (F) Characterization of positive and negative resolution in resins consisting of 0IDX, 14IDX, 18IDX, and 22IDX mixtures. Positive features are indicated with the solid bars and negative features with empty bars. Color of the bar corresponds to energy in mJ·mm^-2^ as shown in the color scale. Error bars represent standard deviation (3 samples printed; 5 measurements per sample). See Tables S3 and S4 for statistical significance. (G) Printing windows of 14IDX and 18IDX mixtures at 16°C and 23°C. (H) Thermal half-life of the DCPI in reference GelMA mixtures with and without IDX, 5–25°C. (I) Branched channel structure increasing from 1mm diameter (bottom) to 750 µm (middle) and 500 µm (end) channels. (J) Gyroid printed in 14IDX mixture. Scale bars 1 mm in B, C, G, H; Scale bars 200 µm in D, E

### Printing of cell-laden structures

Our integrated strategy—combining optical determination, reactivity screening, printability, and analysis—led to the identification of 14IDX as a versatile bioresin with a suitable balance between refractive index matching, photoreactivity, and compatibility with Xolography printing parameters. To validate that this material design strategy translates effectively to cell-laden systems, we investigated the use of the 14IDX formulation to support the fabrication of cell-laden constructs by printing C2C12 myoblast–laden bioresins (5·10^6^ cells·mL^−1^). We considered 5·10^6^ cells·mL^−1^ to be the minimal myoblast concentration required for maturation to myofibers based on previous bioprinting reports.^4,21^

We characterized the printing resolution by fabricating hollow star-shaped structures to assess both positive and negative features (Figure 6A-C). The optimal printing parameters matched those of cell-free prints (2–5 mJ·mm^-2^ with speed 1.0 mm·min^-1^), using a constant gelation and printing temperature of 16 °C. For comparison, the same structures were printed using standard 0IDX bioresins with the same cell concentration (Figure 6D). Similarly to cell-free resins, these required a higher printing energy of 27 mJ·mm^-2^ at a speed of 1.0 mm·min^-1^ to achieve polymerization.

**Figure 6.**
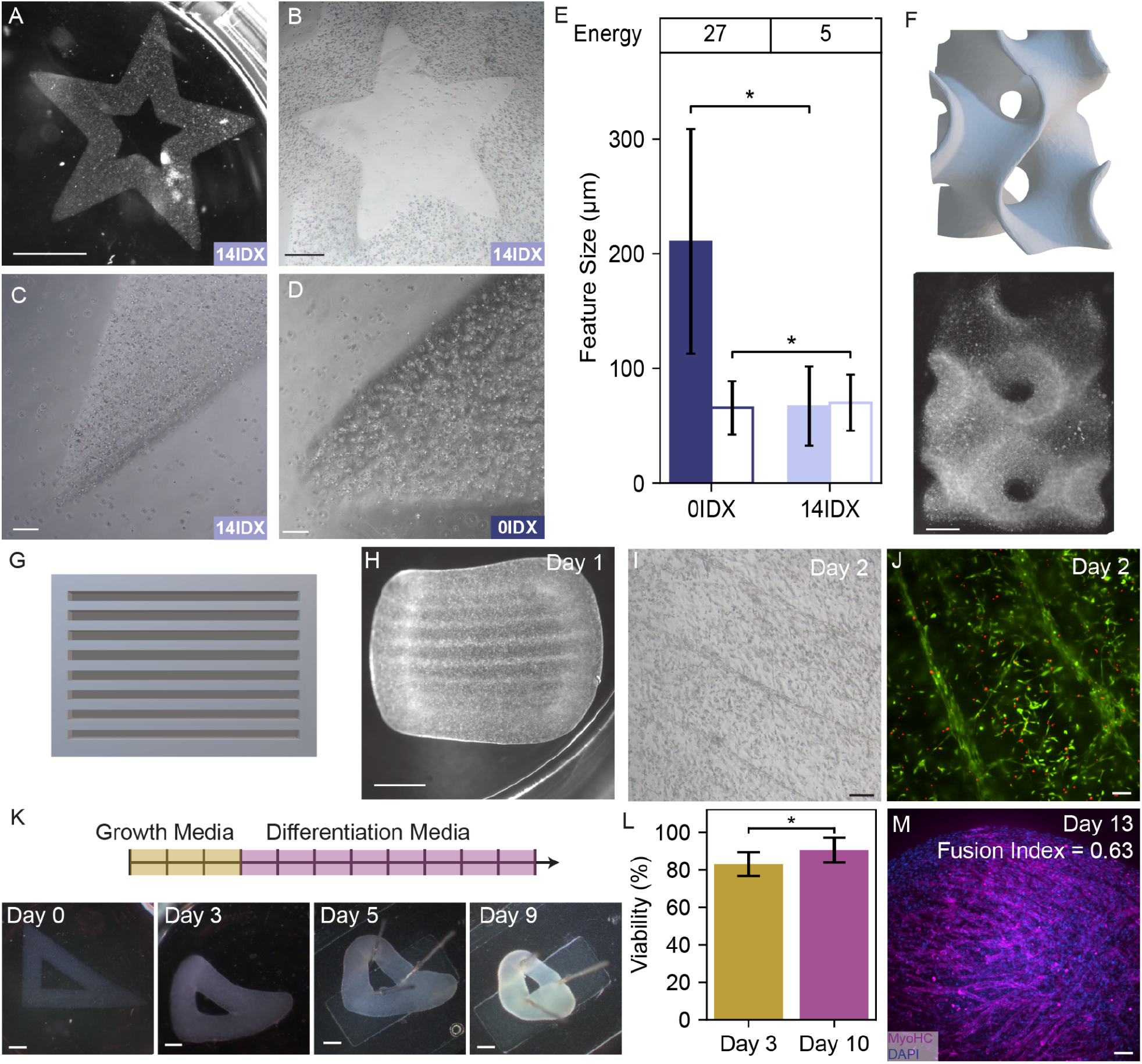
Xolographic printing of high-cell-density structures. (A) Star-shaped construct printed with14IDX bioresin. Close-up of a positive feature (B) and a negative feature (C) in 14IDX prints. (D) Close-up of a positive feature in 0IDX prints. (E) Characterization of feature size in 0IDX and 14IDX bioresins with 5·10^6^ cells·mL^−1^. Positive features are indicated with the solid bars and negative features with the empty bars. The UV energy value (mJ·mm^-2^) for each formulation is shown in the table above. Error bars represent standard deviation (minimum 3 samples printed; 5 measurements per sample). Significant differences (p < 0.01) shown as determined by Welch’s t-test. (F) STL file and stereomicroscope images of a printed gyroid structure. (G) STL file of 300 µm grooved structure used for cell alignment. (H) Stereomicroscope image of grooved structured 24 hours after printing. (I) Inverted microscopy image of cell alignment along groove edges 48 hours after printing. (J) Confocal image of Live/Dead staining 48 hours after printing showing the cell morphology and alignment along groove edges. Live and dead cells are visualized in green and red, respectively. (K) Time course of growth and differentiation of the constructs, showing stereomicroscope images of the constructs. (L) Quantification of cell viability in C2C12-laden constructs at day 3 and day 10 of maturation, based on live/dead staining with nuclear labeling. Viability was calculated as the fraction of dead cells (red-stained nuclei) relative to the total number of cells (n = 4). (M) Confocal image showing C2C12 after day 13 of maturation stained for myosin heavy chain (MyoHC) and nuclei marked with DAPI. Fusion index of this image is 0.63, with an overall sample average (n = 3) of 0.54 ± 0.11. Scale bars 1 mm in A, F, H, K; 200 µm in B, C, D, I; 100 µm in J, M.

The feature size quantification (Figure 6E) showed that 14IDX bioresins achieved positive and negative features in the range of 90 µm and 70 µm respectively, both comparable to the resolution obtained without cells. In contrast, 0IDX prints matched the feature size of their cell-free counterparts but exhibited significantly lower positive resolution than 14IDX. Notably, in both cases we achieved well-defined prints with a lower concentration of BT (450 mM) and higher cell concentration (5·10^6^ mL^-1^) than previously reported (800 mM BT and <1·10^6^ mL^-1^).^22^ However, 24 hours after printing, 0IDX cell prints partially dissolved, indicating that even the higher printing energies failed to achieve complete crosslinking of this formulation. This underscores the improved performance of IDX-containing resins and their suitability for high-cell-density structures. The cell-laden resins maintain high resolution in the z-direction as well, demonstrated by the successful printing of a gyroid with 5·10^6^ cells·mL^-1^ (Figure 6F).

To assess the biocompatibility of the printing process, we evaluated cell viability 48 hours after printing using a Live/Dead assay. Cells remained highly viable (85%) and started spreading within the printed structures, with cells uniformly distributed throughout the printed constructs (Figure S14). We found no difference in cell viability between cells printed in IDX-containing or IDX-free resins, suggesting that the IDX reaction had no observable adverse effect on cell viability (Figure S15). The addition of fibrinogen to the cell-laden resins enhanced cell spreading in the first 48 hours as compared to GelMA-only controls (Figure S16). While GelMA contains arginine-glycine-aspartic acid (RGD) motifs that facilitate cell adhesion, it only partially recapitulates the native extracellular matrix.^55^ Incorporating additional bioactive polymers into GelMA hydrogels is a well-established strategy to create a more cell-supportive environment.^56^ Fibrinogen was chosen as it is widely used in skeletal muscle tissue engineering^57^ and is particularly suitable for mixing with GelMA at 37 °C, as it does not exhibit thermally induced crosslinking.

To further demonstrate that printed architectures can actively influence cell behavior, we fabricated grooved structures designed to guide cellular organization (Figure 6G–H). Inverted microscopy (Figure 6I) and fluorescence imaging following Live/Dead staining (Figure 6J) performed 48 hours after printing revealed that cells aligned along the edges of the patterned structures. This observation highlights the ability of Xolography to encode instructive microarchitectural cues that direct cell organization, a key requirement for applications such as skeletal muscle tissue engineering.

To investigate whether the printed construct environment supports cell self-organization and coordinated differentiation, we cultured printed triangular constructs in growth media for 3 days (Figure 5K) before switching to differentiation conditions for an additional 10 days. Following standard muscle tissue engineering protocol, constructs were tensioned at day 5 to promote maturation. We observed matrix remodeling over the culture period. We evaluated cell viability at day 3 and day 10 of maturation to assess potential long-term cytotoxic effects and observed a slight increase in viability at day 10, consistent with the preceding growth phase (Figure 6L). This confirms that the cells remained viable and did not exhibit any adverse effects throughout the maturation period. Early stages of myogenic differentiation characterized by anisotropically elongated myoblasts and multinucleated cells were confirmed by positive staining for myosin heavy chain (MyoHC, Figure 6M). We characterized the maturation state of the tissue using the fusion index, a well-established metric of myogenic differentiation defined as the proportion of nuclei within MyoHC+ multinucleated myotubes relative to the total number of nuclei.^58^ Using this approach, we measured an average fusion index of 0.54 ± 0.11, consistent with fusion indices reported in engineered muscle tissues.^59,60^ It also suggests that the construct maturation process could be further optimized to result in more mature tissues, as highly differentiated muscle is typically characterized by fusion indices above ∼0.90.^61^ The formation of myotubes in our xolographic constructs confirms that sufficient cell density and biocompatibility were achieved to support myoblast self-assembly and differentiation into multinucleated myotubes, thereby initiating the tissue formation process. Our work marks the first demonstration of cell assembly to tissues in xolographic constructs, as previous studies have only demonstrated sparse cell or spheroid encapsulation.^22–24^ Together, these results demonstrate our platform’s suitability to engineer complex, high-cell-density tissues with biologically relevant features.

## Conclusion

In this work, we reveal the unique dual effect of Iodixanol (IDX) in Bioxolography comprising the combination of increased optical transparency with enhanced reactivity. To date, no systematic study has optimized refractive index-matching under dual-color conditions for cell-laden volumetric printing despite its critical importance for high-cell-density biofabrication. Here, we address this limitation by establishing an IDX-based strategy that restores optical fidelity in highly scattering, cell-rich resins. By tuning resin composition and printing parameters, we leveraged the dual effect of IDX to achieve feature sizes <90 µm and free-standing features down to 100 µm in bioresins containing 5·10^6^ cells·mL^−1^. We developed Xolography resins along an iso-refractive index line, allowing precise control of dual-color reactivity at the matched index for cell-laden prints. Thus, our work provides a validated toolbox for RI-matching in dual-color volumetric printing, enabling constructs with unprecedented optical fidelity, a prerequisite for functional tissue engineering at scale. Moreover, dedicated optimization of printing parameters enabled volumetric printing of hydrogels with complex geometries and tunable mechanical properties. Using skeletal muscle tissue as a model, we demonstrated that refractive index-matched Bioxolography guides cell alignment and supports the formation of mature myofibers in printed muscle tissue constructs, underscoring its potential for functional tissue engineering.

However, specific challenges remain when using IDX in Bioxolography to achieve even higher cell densities. One key aspect relates to the initial excitation of the DCPI at λ_1_ = 375 nm that triggers undesired curing from IDX. While we mitigated this by introducing a hydrogel-compatible radical polymerization inhibitor, red-shifting λ_1_ through molecular DCPI modification alongside an in-depth mechanistic understanding clearly represents the most direct and thus most promising approach to eliminate this side reaction entirely — respective investigations are currently underway. Notably, the reactivity-enhancing effect of IDX appears unique to dual-color photoinitiation systems, highlighting a mechanism-specific phenomenon not observed in conventional light-based printing approaches, in which IDX remains photochemically inert and serves solely for refractive index-matching.

By enabling fine-tuned combinations of reactivity and transparency, Bioxolography expands the design space of cell-laden resins with a material design framework that allows customization to specific cell types and concentrations. We envision that, through its compatibility with complex geometries, multi-material approaches, and potential for high cell densities, this approach will not only enable the engineering of complex, biomimetic tissues, but also open new avenues to engineer and interrogate crosslinking mechanisms in hydrogel systems, ultimately establishing Bioxolography as a foundational technology for next-generation biofabrication and regenerative medicine.

## Experimental Section

### Materials

The solutions for the reference experiments were prepared in DI water, using the following chemicals from given manufacturers without further purification: Iodixanol (BLD Pharm, 99%), PEGDA-575 (Sigma Aldrich, 98%), BT (BLD Pharm, 99%), TEMPOL (Sigma Aldrich, ≥99%), starch (Merck, 99%). For the printing experiments the following substances were used: GelMA (Rousselot, X-Pure 160P80), OptiPrep (StemCell Technologies, Catalog #07820), BT (Sigma Aldrich, >99.0%, #B9754), PBS (Gibco, #10010023), Fibrinogen (Sigma Aldrich, #341573). The used DCPI 5004 was provided by xolo GmbH.

### Instrumentation

Refractive index measurements were performed at a light wavelength of 589 nm and 20 °C (n_D,20_) on a Anton Paar Germany GmbH Abbemat 3200 refractometer. UV-vis absorbance spectra were recorded using an Agilent Cary C60 UV-Vis spectrometer in Brand UV cuvettes (759150 and 759170). All printing experiments were performed on a Xube^2^ volumetric 3D printer (xolo GmbH) in the same cuvettes as used for UV-Vis absorbance measurements.

### Cell Culture

Mouse myoblast cell line C2C12 was obtained from ATCC and cultured using standard protocols. Growth media (GM) consisted of DMEM (Sigma-Aldrich, Catalog # 41966) with 10% fetal bovine serum (Gibco, # A5256801) and 1% Penicillin-Streptomycin (P/S) (Sigma Aldrich, # P0781). Cells were passaged after reaching 80% confluency. Cells from passages 5 to 7 were used for printing experiments. Cell differentiation was induced using differentiation media (DM) consisting of DMEM supplemented with 2% horse serum (Sigma Aldrich, # H1138), 1% P/S and 50 ng·mL^-1^ insulin growth factor (IGF-1) (Thermo Fisher, # 100-11).

### Preparation of bioresin

A 1M BT stock solution in DI water was neutralized to pH 7.4 by addition of aq. HCl and stored at room temperature. Stock solutions of fibrinogen at 80 mg·mL^-1^ fibrinogen in DI water, DCPI 5004 at 6.0 mg·mL^-1^ in DI water, and 0.5 mg·mL^-1^ TEMPOL in DI water were prepared fresh for each printing experiment. To mix 10.0 mL of bioresin, 1.00 g of GelMA was dissolved at 37 °C in a total solvent volume of 9.27 mL by accounting for the volume of dissolved GelMA. The solvent consisted of BisTris, Iodixanol, TEMPOL, DCPI, fibrinogen, and water as necessary to achieve desired concentrations. Final DCPI concentrations were constant at 300 ppm and TEMPOL at 5 ppm for each composition. Fibrinogen was used at 4 mg·mL^-1^. For cell-laden prints, C2C12 cells at a concentration of 5·10^6^ cells·mL^-1^ and a modified nutrient mix were added into the prepared bioresin.^23^

### Cell-free construct processing

All resins were gelled at 16 °C using a Cooling Drybath (ThermoScientific) and printed at 16 °C. Temperature control inside the printing chamber was achieved by the printer’s internal controls and supplemented by an external cooling unit as necessary (see SI section 10 for detailed description). After printing, the cell-free constructs were isolated by washing with warm PBS. Post-crosslinking was performed by incubating the printed constructs with 0.5 U/mL of thrombin in GM for 1 hr.

### Cell-laden construct processing & imaging

Cell-laden constructs were washed into warm GM instead of PBS, and post-crosslinked as described above. The prints were kept in GM for seven days and then moved to DM for an additional ten days. Live/Dead assay (ThermoFisher, #R37601) was performed 48 hours after printing, with Hoechst 33342 (10 ug/mL) for nuclei staining. Averaged values from a minimum of three biological replicates were used for plotting. Constructs were fixed in 4% paraformaldehyde and stained following standard protocols with DAPI, ActinGreen (ThermoFisher, #R37110), and Myosin Heavy Chain (MHC) antibody (ThermoFisher, #50-6503-82). Imaging was performed on a Zeiss Stemi 508 stereomicroscope with GFP module or Nikon Ti2 confocal microscope.

### Image processing

For nuclei counting, images were acquired as z-stacks spanning 100 µm and subsequently maximum intensity projected. The blue channel (Hoechst or DAPI) was intensity-normalized and smoothed using a Gaussian filter. Objects were detected using Laplacian-of-Gaussian (LoG) blob detection. Dead cells (red channel) were identified following intensity normalization and LoG-based object detection. Detection parameters were empirically optimized and verified by visual inspection to ensure accurate identification.

For fusion index analysis, nuclei were detected using the same LoG-based approach. A mask of all detected nuclei was generated, and MyoHC-positive nuclei were manually annotated in Fiji. The fusion index was calculated as the ratio of MyoHC-positive nuclei to the total number of nuclei. Three independent samples were analyzed by quantifying the fusion index in nine regions of interest per sample, derived from three separate maximum intensity projections of 200 µm z-stacks.

### Resolution measurements

All samples were post-cured, stained with trypan blue, and imaged using 4x magnification with an inverted microscope. All five points on the star were imaged, and a minimum of three constructs were prepared for each condition. The resolution was manually measured using ImageJ, and statistical analysis was performed using ANOVA and Tukey’s post-hoc test.

### Mechanical characterization

All samples were equilibrated in PBS for at least 24 hours after printing. Rheological characterization was performed with a viscometer Anton-Paar MC302 (Anton Paar, Graz, Austria) using an 8 mm parallel plate geometry. Measurements were performed on 8 mm diameter hydrogel disks using a strain rate of 0.1% at 1 Hz for 100 s. Averaged data from the last 60 s were used to calculate the storage and loss modulus for each sample. A minimum of three replicates were used per condition.

### Statistical Analysis

Sample numbers are provided in the figure captions, with a minimum of three independent samples for each condition. Error bars in graphs represent standard deviations. For two sample groups, Welch’s independent t-test was used to calculate significance. For multiple group comparisons, ANOVA followed by Tukey’s post-hoc analysis was used.

## Acknowledgments

The work was supported by the SNSF Sinergia Grant No. 216727. At ETH Zurich, this work was done within the framework of the ALIVE initiative (Advanced Engineering with Living Materials) and funded by the SFA-AM program (Strategic Focus Area – Advanced Manufacturing). N.F.K. acknowledges funding from the Federal Ministry of Economic Affairs and Energy (BMWE, grant No. 50WM2446). A. Balciunaite, A. Badolato and R.K.K. thank Dr. Costanza Giampietro and Prof. Dr. Edoardo Mazza for the support and use of the confocal microscope and Prof. Dr. Mark Tibbitt and his group members for the support and use of the rheometer. S.H. thanks the Einstein Foundation Berlin as well as Humboldt University for generous support.

