## Supplemental Information for "Cell-Dense Bioink Design for Xolography: Coupling Refractive Index-Matching with Increased Photoreactivity"

### Table of Contents

1) Custom-built transmittance setup 1
2) Transmittance measurements 1
3) Extinction coefficient of IDX 2
4) Calculated light sheet intensity profiles 3
5) Reference system experiments for dual-color reactivity gain 4
6) Mechanistic studies on the influence of IDX on dual color photopolymerization 5
7) Radical polymerisation inhibitor screening 8
8) Printing examples for 22IDX composition 8
9) Mechanical Characterization 9
10) Printing Methodology 10
11) Biological characterization 11
12) References 11

### 1) Custom-built transmittance setup

The setup comprises two LED light sources, one with narrow emission with a central wavelength of 375 nm in the UV regime (365-385 nm, bandwidth: 9 nm), representing the UV light sheet and one in the vis regime (500-750 nm, bandwidth: 104 nm) with a central wavelength of 565 nm, which corresponds to the broadband vis projection of the printer. After merging of both light paths by a dichroic mirror and collimation, the dual-color beam is passed through the sample before being analyzed in intensity by a power meter with a background-corrected thermal sensor. The spectrometer was checked for functionality by measuring different ND filters with known degrees of transmittance. The sample is placed between the collimation optics and the detector in a 3D printed cuvette holder containing further iris shades for light-beam modulation, ensuring that only direct transmitted light is detected. As sample containers, small-volume variants (1.0 mL) of the cuvettes used for actual printing (3.5 mL) were used to ensure comparability. For illumination, mounted LEDs (M375L4, M565L3) were used combined by a DMLP425R mirror and collimated with ACL2520U optics. The output power was measured with a PM16-401 USB power meter. All named components were purchased from Thorlabs.

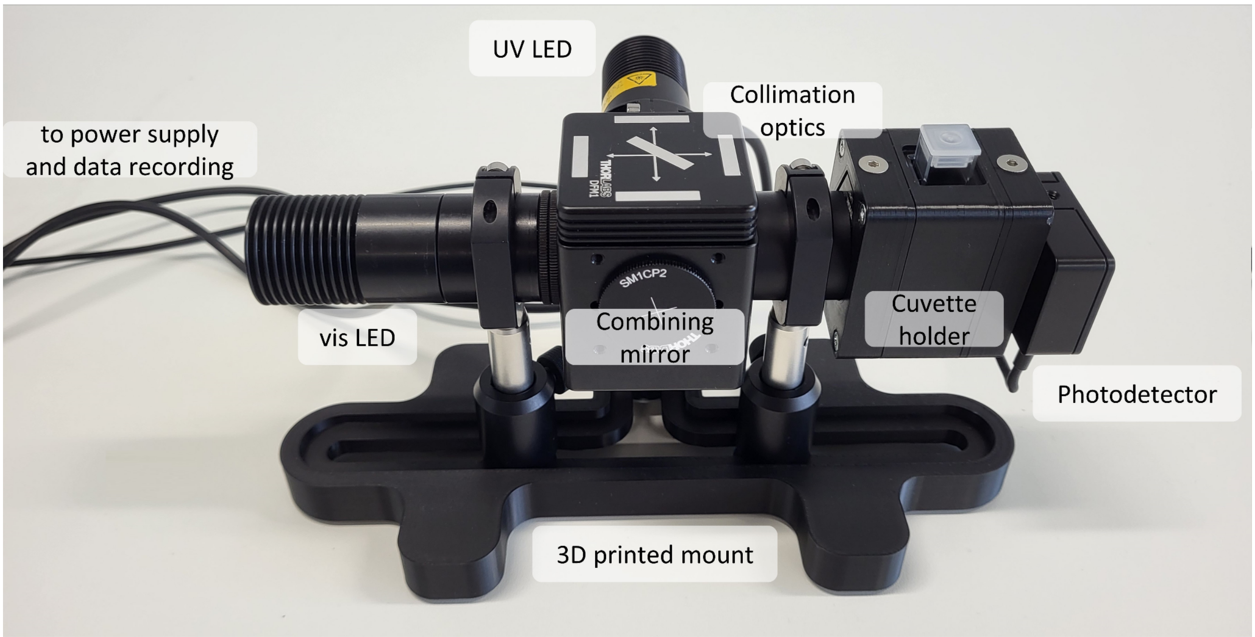


**Figure S1.** Photography of the developed dual-color transmittance setup for both the UV- and the visible light-range.

### 2) Transmittance measurements

For the transmittance measurements shown in the main text Figure 2, C2C12 cells were suspended in GelMA (10% w/v in water, 186 mM BT). The required IDX amounts were calculated in a DoE-matrix. The reference samples contained each component except the cell suspension; the missing volume was balanced with PBS solution.

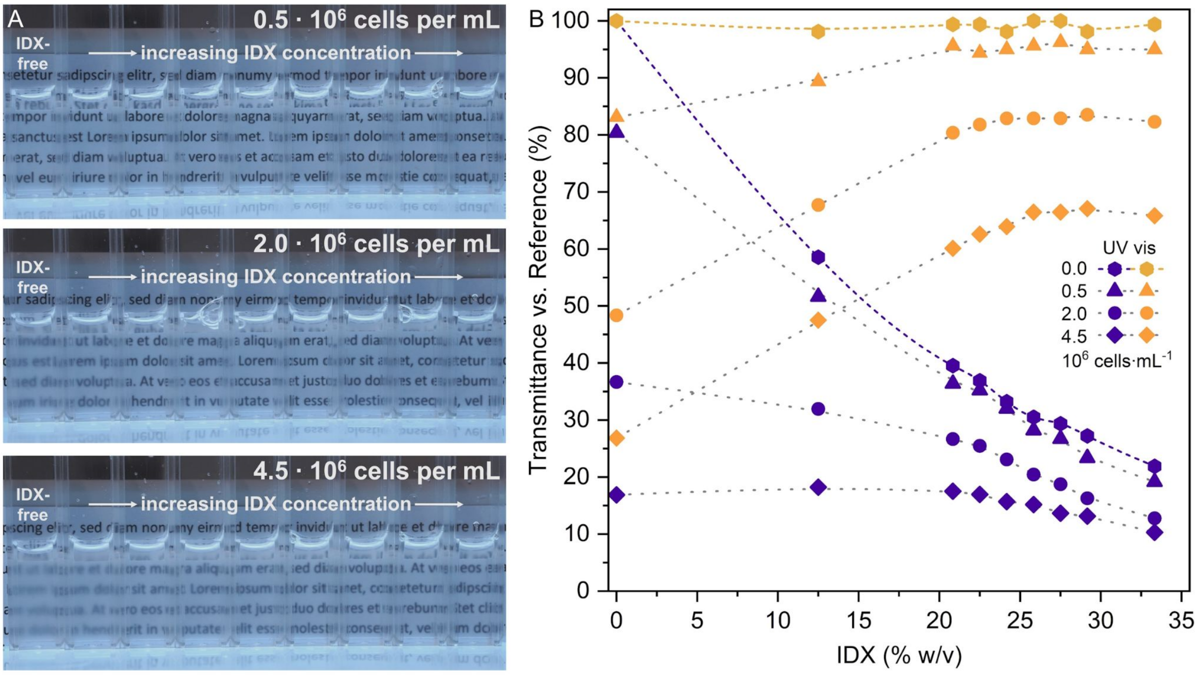


**Figure S2.** Photographs (A) and transmittance analysis (B) of cell suspensions in water, showcasing that for higher cell densities, even the 'matched' conditions do not lead to a complete transparency. Simultaneously, the UV light transmittance is strongly attenuated, and therefore limiting the usable amount of cells and IDX.

### 3) Extinction coefficient of IDX

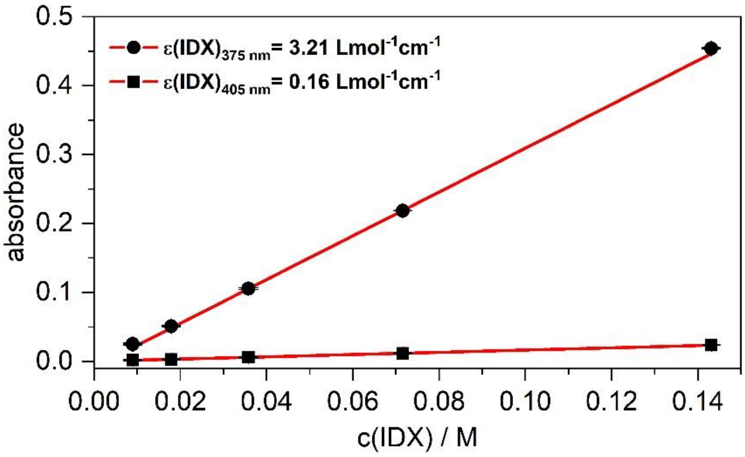


**Figure S3.** Determination of the extinction coefficient of IDX in water for 375 nm (circles) and 405 nm (boxes).

### 4) Calculated light sheet intensity profiles

As described in the original Xolography paper, the idealized intensity profiles I(x) of the light sheet across the cuvette width x can be calculated using a simplified model based on the Lambert-Beer-Bouguer law:[S1]

$$I(x) = \frac{I_0}{2} \times \left[ e^{-\left(\frac{OD^{1cm}}{1cm} + \alpha_S\right)x} + e^{-\left(\frac{OD^{1cm}}{1cm} + \alpha_S\right)(1-x)} \right]$$

By applying this model to different scattering coefficients and using the measured transmittance of IDX (18% w/v corresponds to OD = 0.37 at 375 nm), it is found that illuminating from both sides of the cuvette produces a more homogeneous intensity distribution across the width (Figure S4A and B). However, as the scattering coefficient increases as expected for cell-laden mixtures, this homogeneity is significantly reduced, highlighting the need for refractive index matching to mitigate scattering. This conclusion holds for both UV and visible light. However, at longer wavelengths such as 405 nm, the intensity distribution is inherently more uniform (Figure S4C), due to the lower absorbance of IDX at this wavelength (OD = 0.02 at 405 nm for 18% w/v). Additionally, as the scattering coefficient approximately scales with the inverse fourth power of the wavelength ($\alpha_S \propto \lambda^{-4}$), scattering is intrinsically lower at 405 nm than at 375 nm. In combination, these effects underscore the importance of developing efficient DCPIs that are switchable at 405 nm (or higher) instead of 375 nm, to reduce absorption and scattering and thereby improve light-sheet homogeneity in refractive index matched Xolography using IDX.

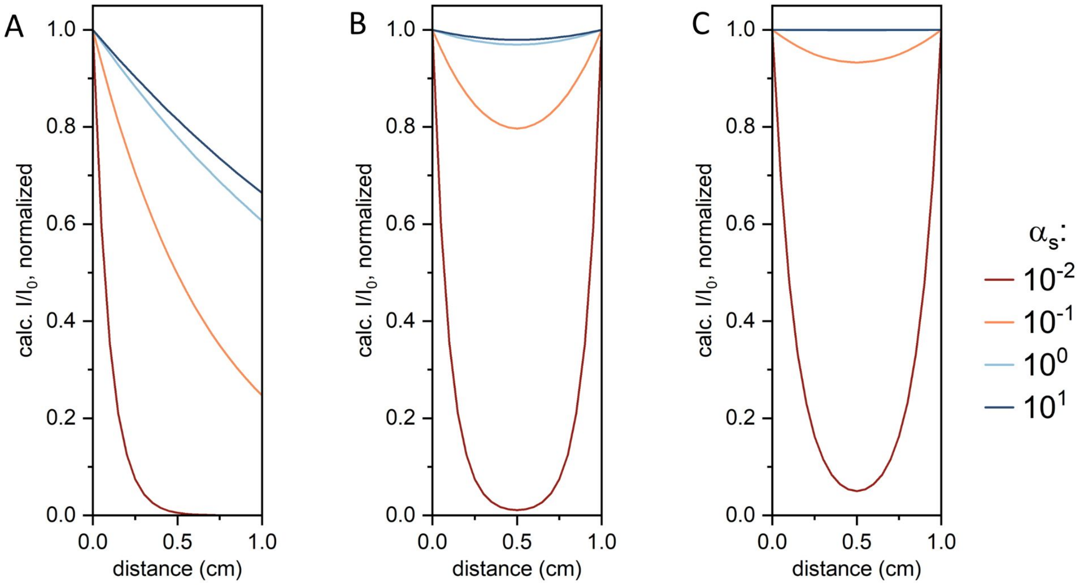


**Figure S4.** Calculated light sheet intensity profiles for IDX-containing solutions at varying scattering coefficients $\alpha_S$, A: unidirectional illumination at 375 nm (OD = 0.37), B: dual-sided illumination at 375 nm (OD = 0.37), which produces a more uniform intensity distribution across the cuvette width. The results demonstrate that bidirectional illumination improves homogeneity; however, the degree of uniformity strongly depends on the magnitude of the scattering coefficient, highlighting the need to reduce scattering throughout the volume. C: Dual-sided illumination at 405 nm, where both the optical density (OD = 0.02) and the scattering coefficient are reduced due to the lower absorption of IDX and the wavelength dependence ($\alpha_S \propto \lambda^{-4}$), resulting in an improved intensity uniformity.

### 5) Reference system experiments for dual-color reactivity gain

The dual-color reactivity gain (DCRG) percentage is calculated as:

$$\% \text{ DCRG} = \frac{t_{gel}\ (\text{UV}-\text{only}) - t_{gel}\ (\text{UV}+\text{vis})}{t_{gel}\ (\text{UV}-\text{only})} \times 100$$

with $t_{gel}$ given in Tables S1 and S2.

The measured light intensities for all experiments were 21 mW•cm$^{-2}$ in the case of 375 nm and 120 mW•cm$^{-2}$ in the case of 565 nm illumination.

**Table S1.** Gelation times for PEGDA575-solutions (20% v/v) in water with varying IDX and BT contents along the iso-RI line.

| | Substance | | Gelation time $t_{gel}$ /s | | DCRG /% |
|---|---|---|---|---|---|
| | BT /mM | IDX /% w/v | UV-only | UV+vis | $\frac{t_{gel}(\text{UV-only})-t_{gel}(\text{UV+vis})}{t_{gel}(\text{UV+vis})}$ |
| **1** | 500 | 12.9 | 84 ± 2 | 58 ± 0 | 44.0 |
| **2** | 450 | 14.0 | 84 ± 2 | 60 ± 1 | 40.3 |
| **3** | 400 | 15.0 | 88 ± 1 | 63 ± 0 | 38.9 |
| **4** | 350 | 16.0 | 92 ± 4 | 67 ± 1 | 37.6 |
| **5** | 300 | 17.0 | 95 ± 2 | 70 ± 2 | 36.0 |
| **6** | 250 | 18.0 | 100 ± 2 | 78 ± 1 | 29.7 |
| **7** | 200 | 19.1 | 113 ± 3 | 91 ± 2 | 23.6 |
| **8** | 150 | 20.1 | 130 ± 3 | 110 ± 3 | 18.7 |
| **9** | 100 | 21.1 | 165 ± 2 | 144 ± 2 | 15.0 |
| **10** | 50 | 22.1 | 229 ± 6 | 206 ± 5 | 11.4 |
| **11** | 0 | 23.2 | 924 ± 6 | 745 ± 3 | 24.0 |

**Table S2.** Gelation times for PEGDA575-solutions (20% w/v) in water with varying compositions of DCPI (300 ppm) / BT (250 mM) / IDX (18% w/v) for UV only and UV+vis illumination.

| | Substance | | | Gelation time $t_{gel}$ /s | | DCRG /% |
|---|---|---|---|---|---|---|
| | DCPI | BT | IDX | UV-only | UV+vis | $\frac{t_{gel}(\text{UV-only})-t_{gel}(\text{UV+vis})}{t_{gel}(\text{UV+vis})}$ |
| A | ✓ | ✓ | ✗ | 280 ± 1 | 184 ± 1 | 51.9 |
| B | ✓ | ✓ | ✓ | 102 ± 5 | 78 ± 1 | 31.0 |
| C | ✗ | ✓ | ✓ | 99 ± 1 | 101 ± 1 | <0.0 |
| D | ✗ | ✗ | ✓ | > 1800 | > 1800 | - |
| E | ✓ | ✗ | ✓ | > 1800 | > 1800 | - |
| F | ✗ | ✓ | ✗ | no gelation | no gelation | - |

### 6) Mechanistic studies on the influence of IDX on dual color photopolymerization

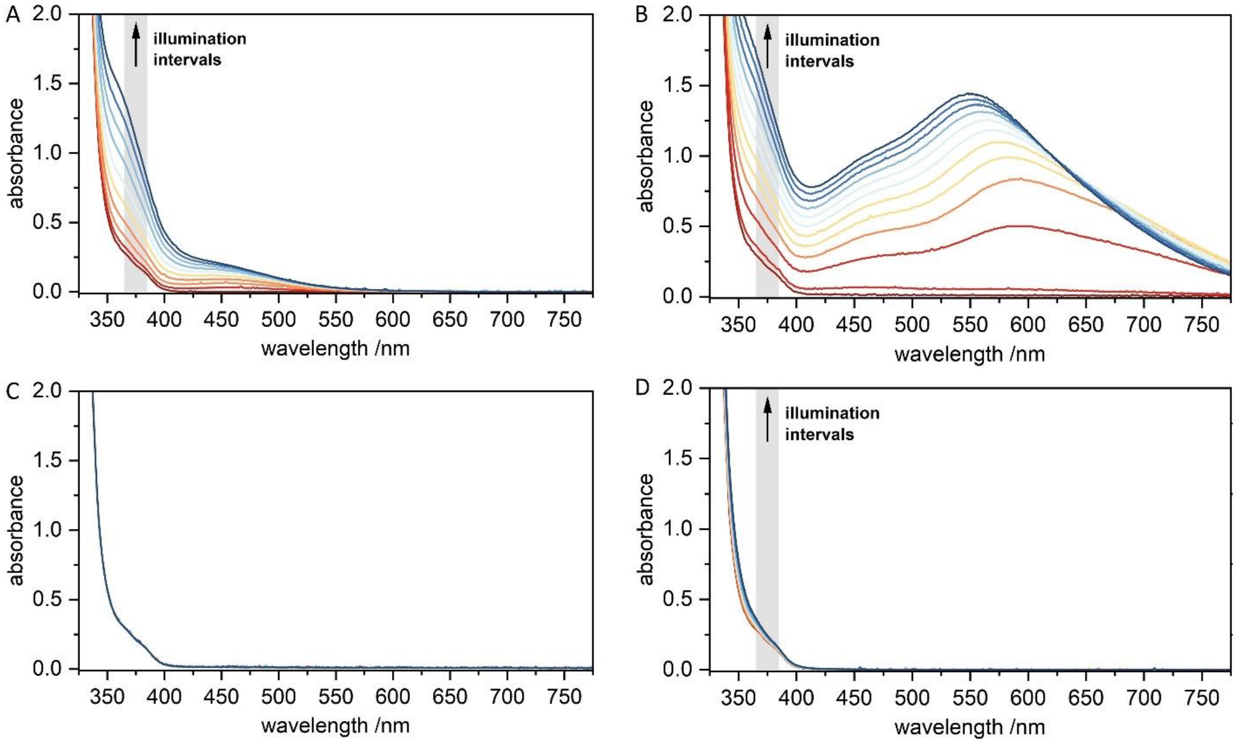


**Figure S5.** Verification for the release of molecular iodine from IDX under 375 nm illumination (grey box). A: Formation of the broad absorbance band of iodine in water. B: Formation of the characteristic blue colored complex after addition of conc. starch solution to the IDX solution and illumination. C: No band formation for same sample as B but without illumination (negative control). D: In the presence of BT, the yellow band of formed $I_2$ is bleached due to the reduction from $I_2$ to $I^-$ in the presence of tertiary amines.

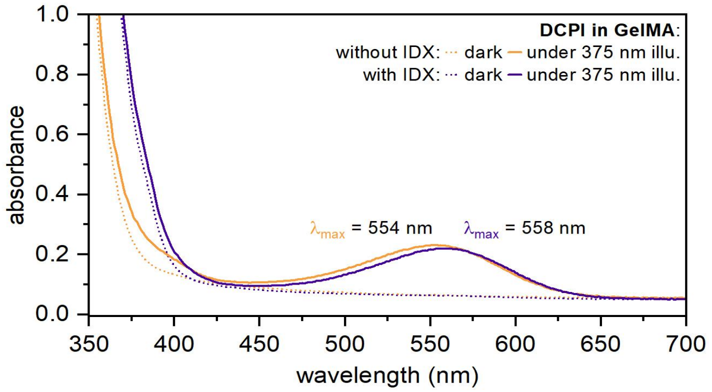


**Figure S6.** UV Vis absorbance spectra of both DCPI 5004 forms (SP dotted, MC solid) in the presence (purple) and absence (orange) of IDX in the GelMA matrix (10% w/v + 250 mM BT, 15 °C). The MC spectra were recorded after 375 nm irradiation.

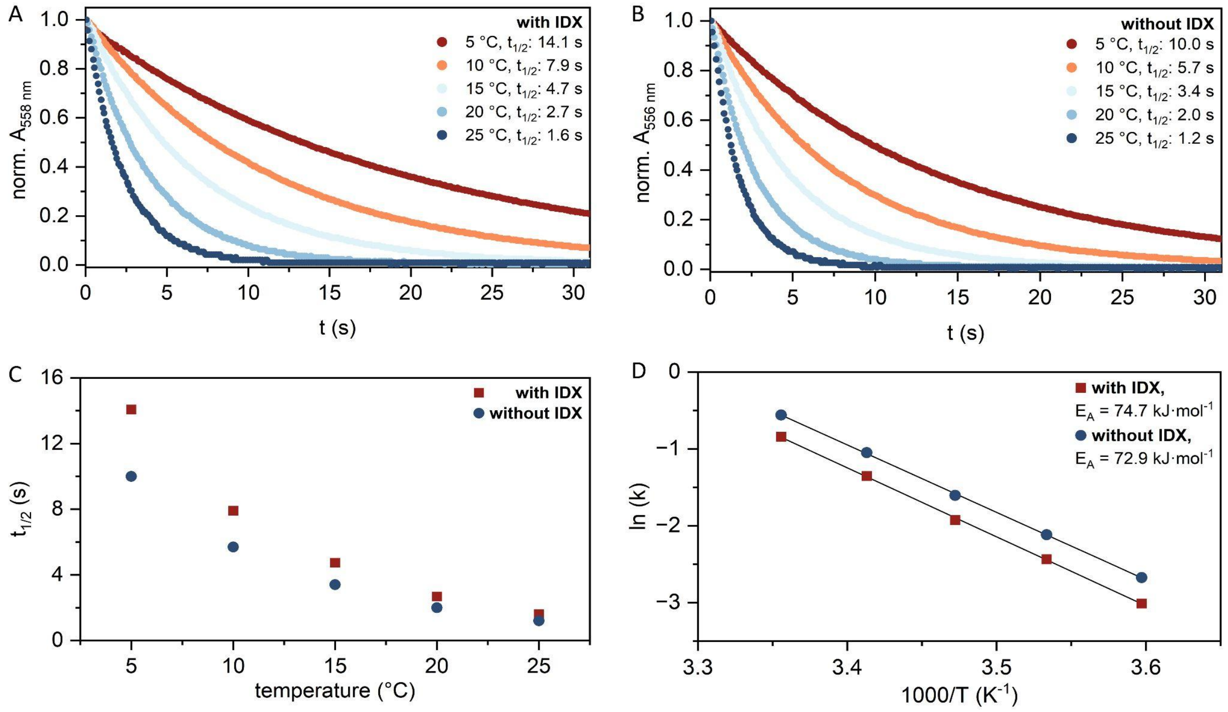


**Figure S7.** A+B: Development of visible light absorbance band at $\lambda_{max}$ for the MC form of DCPI5004 (558 nm with IDX, 554 nm without IDX). C: Determination of the thermal half-life and thermal activation barriers of the MC form based on A+B at different temperatures, as described in references.[S1, S2]

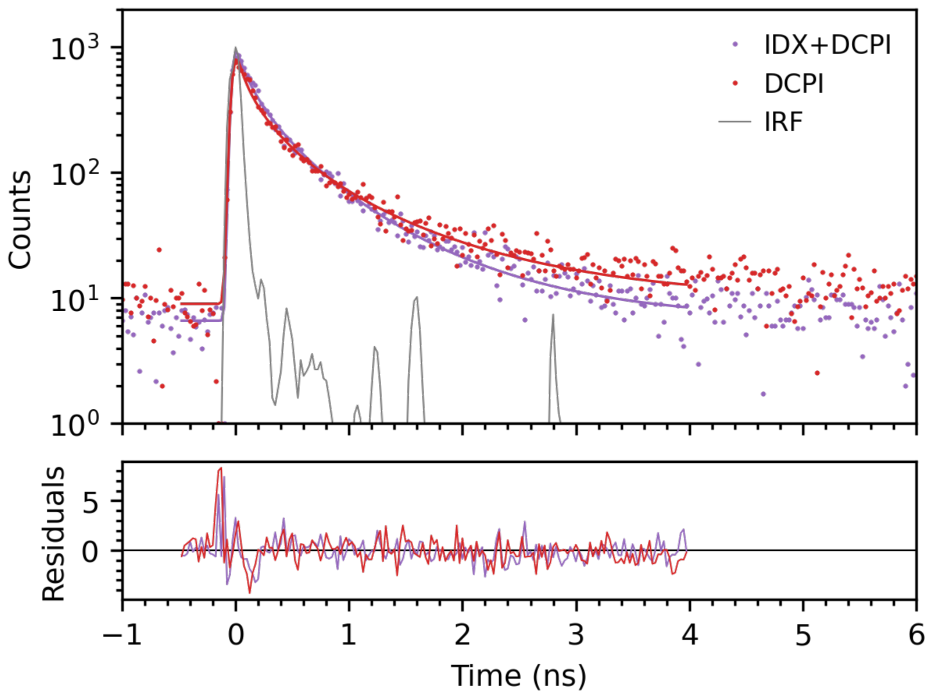


**Figure S8.** Time-correlated single photon counting (TCSPC) measurements of the excited-state lifetime of the merocyanine form (MC*) of DCPI in GelMA at the photostationary state (PSS). Samples were excited at 590 nm and emission was detected at 660 nm. The fluorescence decays were fitted using a stretched exponential function convoluted with a Gaussian instrument response function (IRF). In the presence of IDX, the fluorescence lifetime decreases from 0.682 ns to 0.507 ns, indicating accelerated excited-state deactivation.

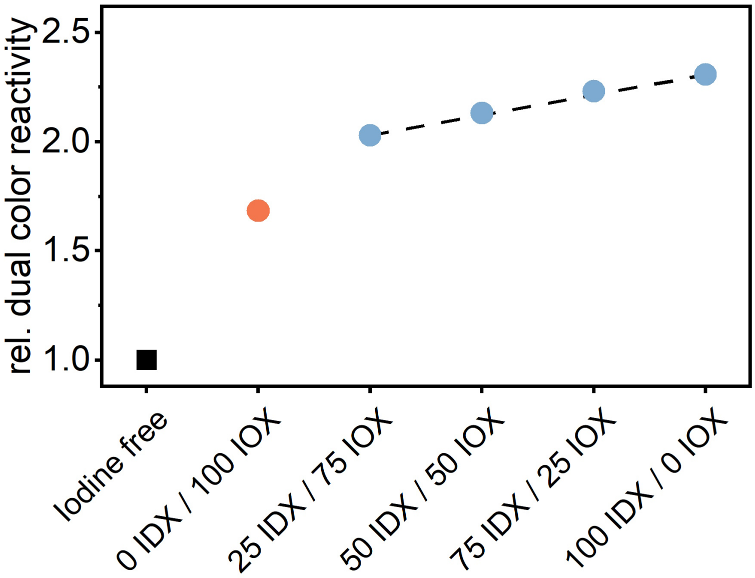


**Figure S9.** Relative dual color reactivity of the PEGDA reference system (20 %w/v PEGDA, 450 mM BT, 300 ppm DCPI). The iodixanol-containing formulation 100 IDX / 0 IOX (14.00 % w/v, corresponding to 68.7 mg mL$^{-1}$ iodine) exhibits the highest relative reactivity (blue point). Systematic substitution of iodixanol with iohexol at constant total iodine concentration leads to a gradual decrease in reactivity. With increasing iohexol fraction, the reactivity decreases approximately linearly, reaching lower values for the iohexol-only sample (14.83 % w/v, 68.8 mg mL$^{-1}$ iodine; orange point), which still remains above the iodine-free reference (black square). The offset between the iohexol-only sample and the iodixanol-containing series indicates that iodixanol introduces an additional contribution beyond iodine concentration alone, consistent with additive-specific interactions such as local environment or microviscosity effects.

### 7) Radical polymerization inhibitor screening

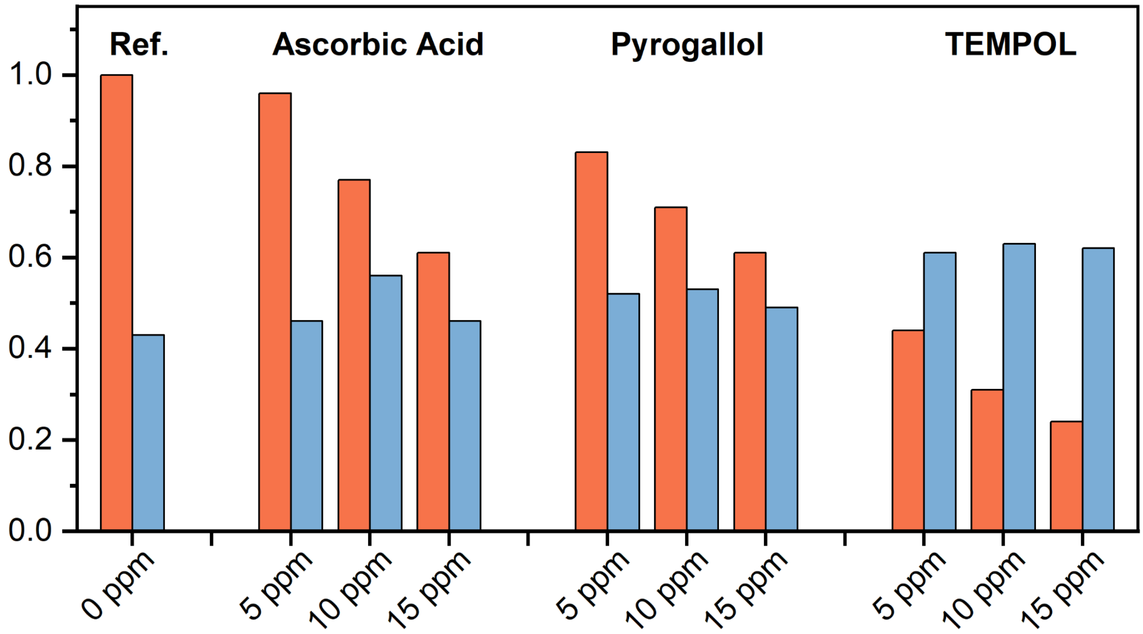


**Figure S10.** Relative overall dual color reactivity (red) and DCRG (blue) of mixtures with different concentrations of the tested inhibitor ascorbic acid, pyrogallol and TEMPOL against an inhibitor free reference, indicating the highest inhibition strength of TEMPOL.

### 8) Printing examples for 22IDX composition

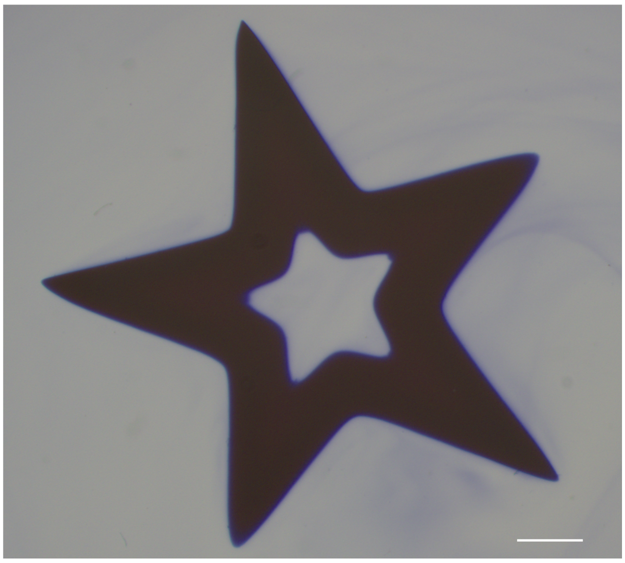

**Figure S11.** Examples of 22IDX mixture printed using E = 13 mJ•mm$^{-2}$ and s = 1.0 mm•mm$^{-1}$. The star is slightly asymmetric, as noticeable from the different tip sizes. Scale bar 1 mm.

### 9) Mechanical Characterization

**Table S3**: Tukey's post-hoc results for positive resolution (corresponding to Figure 5F).

| group1 | group2 | meandiff | p-adj | reject |
|---|---|---|---|---|
| idx_0 | idx_14 | -143.4845 | 0 | True |
| idx_0 | idx_18 | -130.7863 | 0 | True |
| idx_0 | idx_22 | -124.835 | 0 | True |
| idx_14 | idx_18 | 12.6983 | 0.9446 | False |
| idx_14 | idx_22 | 18.6495 | 0.8465 | False |
| idx_18 | idx_22 | 5.9512 | 0.9937 | False |

**Table S4**: Tukey's post-hoc results for negative resolution (corresponding to Figure 5F).

| group1 | group2 | meandiff | p-adj | reject |
|---|---|---|---|---|
| idx_0 | idx_14 | 4.5828 | 0.9803 | False |
| idx_0 | idx_18 | 44.595 | 0.0022 | True |
| idx_0 | idx_22 | 109.4813 | 0 | True |
| idx_14 | idx_18 | 40.0122 | 0.0132 | True |
| idx_14 | idx_22 | 104.8985 | 0 | True |
| idx_18 | idx_22 | 64.8863 | 0 | True |

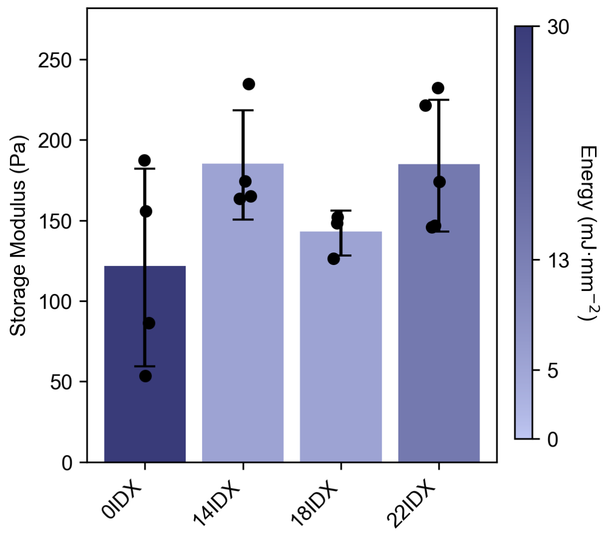


**Figure S12.** Storage modulus comparison of hydrogels printed with speed of 1 mm/min and no post-curing.

### 10) Printing Methodology

**Description of printing setup**: We used a Xube² printer that was connected to an external AC unit (AEG AXP26). This was a temporary fix that was implemented due to time constraints due to a faulty overheating sensor in the Peltier unit in our Xube². The cold air of the AC unit was connected to the air inflow of the printing chamber. We reached a stable 16 °C in the printing chamber within 15 minutes with this method, and were able to keep the temperature at 16 °C by controlling the external AC unit. The Xube² contains an internal temperature control that should be capable of maintaining 16 °C in the printing chamber. However, all experiments in this manuscript were completed using the described setup.

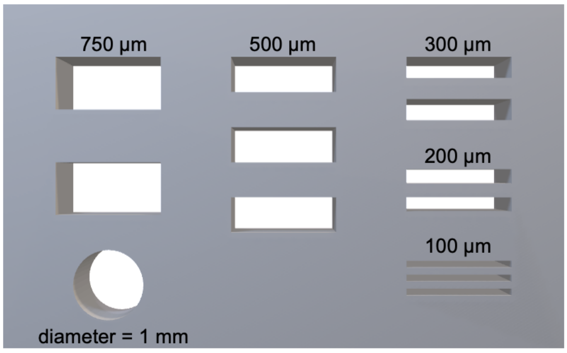


**Figure S13.** STL file of the resolution grid used to determine the printing windows.

### 11) Biological characterization

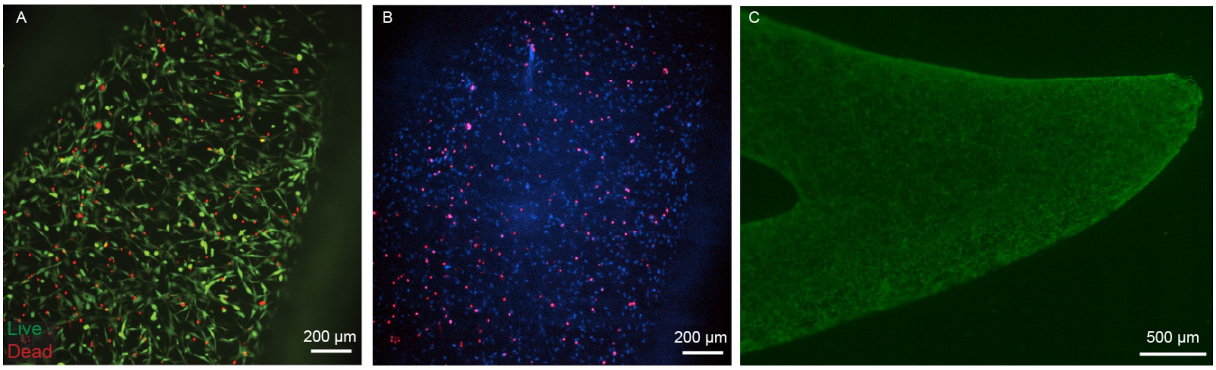


**Figure S14.** (A) Representative confocal microscopy image of constructs stained for live/dead viability 48 hours post-printing. (B) Quantitative analysis indicated ~85% viable cells, calculated based on the fraction of dead cells (red-stained nuclei) relative to the total number of nuclei (blue). (B) Representative stereomicroscope image illustrating the homogeneous distribution of viable cells within the printed construct 48 hours post-printing.

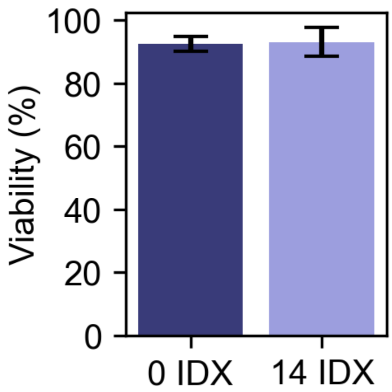


**Figure S15.** Cell viability of printed constructs in the presence and absence of iodixanol. Bar graph showing the percentage of viable cells in constructs printed using either iodixanol-containing resin (14IDX) or with no iodixanol (0IDX). Quantification at day 3 post-printing demonstrated high cell viability in both conditions, with no statistically significant difference observed between groups (n = 3 - 4, Student's t-test p = 0.6)

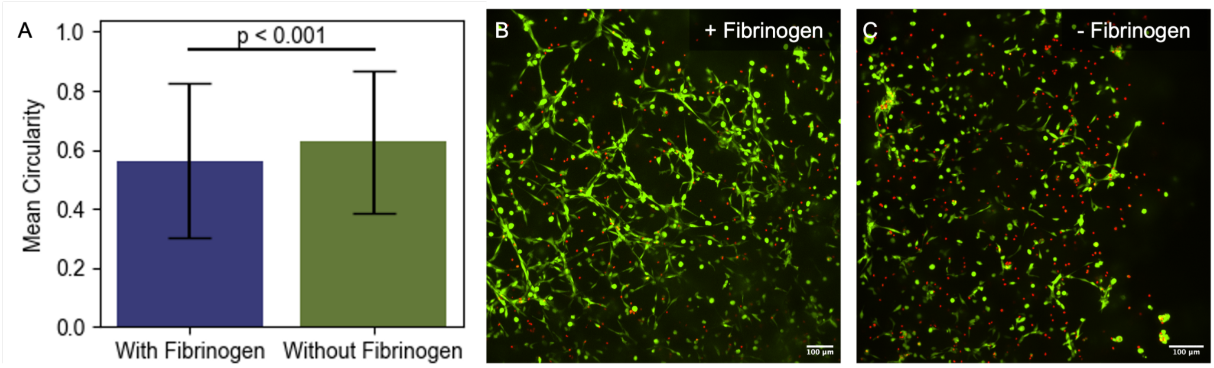


**Figure S16.** Influence of fibrinogen in printing mixture of cell spreading 48 hours after printing. (A) Quantitative analysis of cell circularity in constructs printed with and without fibrinogen, done with FIJI. (B) Representative live/dead staining images of cells within printed constructs, comparing conditions with and without fibrinogen.
